# Contrasting trends in forest growth and mortality of major European tree species under increasing climatic stress

**DOI:** 10.64898/2026.05.18.725878

**Authors:** Miriam Bravo-Hernández, Julen Astigarraga, Susanne Suvanto, Cristina Grajera-Antolín, Marta Rodríguez-Rey, Albert Vilà-Cabrera, Thomas A.M. Pugh, Miguel A. Zavala, Adriane Esquivel-Muelbert, Julián Tijerín-Triviño, Lorena Gómez-Aparicio, Julien Barrere, Verónica Cruz-Alonso, Jonas Fridman, Georges Kunstler, Andrzej Talarczyk, Mart-Jan Schelhaas, Sara Villén-Pérez, Paloma Ruiz-Benito

**Affiliations:** Universidad de Alcalá, Madrid, Spain; Lund University, Lund, Sweden; Luke - Natural Resources Institute Finland, Helsinki, Finland; University of Vic – Central University of Catalonia, Barcelona, Spain; University of Birmingham, Birmingham, UK; IRNAS – Instituto de Recursos Naturales y Agrobiología de Sevilla, Seville, Spain; Université Grenoble Alpes, INRAE LESSEM, Saint-Martin-d’Hères, France; Universidad Complutense de Madrid, Madrid, Spain; Swedish University of Agricultural Sciences, Umeå, Sweden; Forest and Natural Resources Research Centre Foundation, Warsaw, Poland; WERN - Wageningen Environmental Research, Wageningen, Netherlands

**Keywords:** mortality occurrence, mortality intensity, vapor pressure deficit, climatic edge, stand development, needle-leaved, broad-leaved, National Forest Inventories

## Abstract

Forests play a crucial role in mitigating climate change as primary terrestrial carbon sinks. While some studies suggest that global warming enhances forest productivity, a growing body of evidence highlights detrimental impact primarily driven by increased water stress. Yet the extent to which positive effects of climate change offset its negative impacts on tree species productivity remains unclear at large spatial extents. We assessed forest growth and mortality for the 21 most abundant tree species in Europe using National Forest Inventory data from more than 50,000 plots and 700,000 trees to disentangle the relative importance of climate and forest structure. Specifically, we examined how vapor pressure deficit (VPD) anomalies across species’ climatic edges and stand developmental stages affect forest growth and mortality occurrence and intensity (i.e. whether mortality occurred and the amount of basal area lost). Then, we aggregated the responses across species and separately for broad-leaved and needle-leaved species to assess whether forest growth and mortality differed between major functional groups. Although the importance of forest growth and mortality drivers varied markedly among species, climate had a stronger influence on mortality than on growth, particularly in needle-leaved species. Forest growth declined and mortality increased along VPD anomaly in most species and forests studied. Responses were most pronounced at arid species’ edges in early-stage broad-leaved forests and at wet edges in late-stage needle-leaved forests, where differences between functional groups were also highest. We evidence the need to parametrise species-specific models of forest growth and mortality across large spatial extents to better understand and predict effects of climate change on forest productivity. In addition, our results emphasize the importance of improving the understanding of forest mortality processes given the strong influence of climate on mortality, while also further studying vulnerable populations to climate change in arid edges of species distributions.

## 1. Introduction

Forests are acting as global carbon sinks, absorbing roughly half of the fossil fuel emissions over the last three decades, and therefore, constitute an essential ecosystem for climate change mitigation (FAO 2020; Pan et al. 2024). Understanding how forest carbon dynamics are impacted by climate change is key, yet it remains a subject of debate (Reyer et al. 2014; Sperry et al. 2019; McDowell et al. 2020). Multiple studies suggest increased forest productivity due to the positive effects of climate warming and CO_2_ fertilisation (Song et al. 2022; Norby 2025). However, climate change-related disturbances such as heatwaves, droughts or insect outbreaks can decrease forest performance (Reyer et al. 2017; Tijerín-Triviño et al. 2025). Some evidence suggests that the negative effects of climate change on forest productivity may offset its positive effects (Yuan et al. 2019; Hogan et al. 2024), although it depends on the biogeographical region (Ruiz-Benito et al. 2014). Furthermore, climate effects may differ between growth and mortality processes (Kunstler et al. 2021; Evans et al. 2025) and among species and functional groups (Astigarraga et al. 2024; Idoate-Lacasia et al. 2025). Nonetheless, large-scale evaluations simultaneously assessing growth and mortality responses across multiple species remain scarce (but see Anderegg et al. 2020; Kunstler et al. 2021; Dupont-Leduc et al. 2024), limiting our understanding of how climate change affects forest productivity across broad environmental gradients.

The effects of climate change on vegetation are well documented (Evans et al. 2025), but forest susceptibility is also dependent on other factors such as water availability and stand characteristics (Hampe & Petit 2005; McDowell et al. 2020). It is well known that warming directly increases atmospheric dryness, which can lead to reduced forest productivity (McDowell et al. 2020), with populations growing at the arid edge of the species distributions potentially being more susceptible to climate change, although they might also be better adapted to water stress than populations in wetter regions (Isaac-Renton et al. 2018; Ghouil et al. 2020). Identifying the populations most vulnerable to climate change and their locations within the species distribution remains complex and debated, highlighting the need for large-scale analyses that capture the high climatic heterogeneity within species’ distributions (Kunstler et al. 2021). Stand development can also influence forest productivity in interaction with climate (Astigarraga et al. 2024; Lunde et al. 2025). In early successional stands, where tree competition is higher and the climatic buffering capacity of canopy dominant trees is weak, individuals located at the species driest edge could experience the greatest productivity declines (van Breugel et al. 2012; Máliš et al. 2023). The consequences could be particularly important in Northern Hemisphere forests where management has strongly altered stand age, size distributions, and density (Pugh et al. 2024; Lunde et al. 2025). In Europe, most forests are at early stages of stand development due to intensive and recurrent past management, with managed areas being often younger, denser, and structurally more homogeneous than unmanaged forests (e.g. Vilén et al. 2012; McGrath et al. 2015; Ruiz-Benito et al. 2012). Consistently, managed forests may experience greater impacts from warming and water-stress (Vilà-Cabrera et al. 2023). Reducing uncertainty about which forests are the most susceptible to climate change is therefore crucial for informing conservation and management strategies. In this context, extensive forest inventories offer a powerful tool for assessing broad-scale forest responses due to their large spatial coverage and high-quality data (Ruiz-Benito et al. 2020; International Forest Mortality Network 2025).

Numerous studies have quantified changes in productivity dynamics in response to climate change using forest inventory data, providing broad insights into productivity trends and drivers across large spatial extents (Schelhaas et al. 2018; Hogan et al. 2024; Tijerín-Triviño et al. 2025). However, understanding the underlying demographic processes driving these changes remains a challenge (McDowell et al. 2020). Forest productivity is determined not only by a single productivity component such as forest growth, often used as a proxy for productivity, but by the balance between productivity gains (forest growth and recruitment) and losses (forest mortality), which arises from the demographic dynamics of individual trees (Stephenson & van Mantgem 2005; Benito-Garzón et al. 2013). Despite the contrasting contribution of forest growth and mortality to forest productivity, and the numerous studies that have examined either process individually (Pretzsch et al. 2014; Senf et al. 2018; Yuan et al. 2019; Idoate-Lacasia et al. 2025), few studies have assessed their relative contribution to productivity at large spatial extents (but see Wunder et al. 2008; Ruiz-Benito et al. 2017a; Rozendaal et al. 2020). Moreover, how climate change influences both processes simultaneously is rarely assessed at large spatial extents (Anderegg et al. 2020; Kunstler et al. 2021; Dupont-Leduc et al. 2024). Forest growth is expected to be the main contributor to productivity, especially at early stages of development (Dupont-Leduc et al. 2024), partly due to the stochastic nature of mortality (Franklin et al. 1987). However, when mortality happens, it can strongly impact forest productivity, removing a large amount of biomass (Franklin et al. 1987; Stephenson et al. 2014) and potentially offsetting the biomass added by growth and recruitment (Bauman et al. 2022). Understanding how both growth and mortality respond to climate change is therefore essential for accurately assessing forest productivity under a changing climate and understand the outcomes of their balance.

Responses to climate change might vary significantly among species, since different species have different characteristics and strategies to cope with climate (Choat et al. 2018). Several authors have found contrasting species-specific responses to climate change (Benito-Garzón et al. 2013; Changenet et al. 2021; Astigarraga et al. 2024), highlighting the need to consider the species-level when evaluating forest dynamics under climate change (Ruiz-Benito et al. 2013; Neupane et al. 2023), since they encapsulate species-specific traits that translate into particular life strategies (Díaz et al. 2016). However, different species can share certain functional traits, which might lead to similar responses in species with similar life strategies or evolutionary histories (Fei et al. 2017). For instance, broad-leaved and needle-leaved species have exhibited divergent responses to climate change (Fei et al. 2017; Zhao et al. 2024), with needle-leaved species showing particularly high susceptibility to climate change (Carnicer et al. 2013; Sáenz-Romero et al. 2019; Qi et al. 2023). The heightened susceptibility of needle-leaved species has been linked to their conservative water-use strategy, characterised by early stomatal closure to prevent hydraulic failure (McDowell et al. 2008). While effective in the short-term, this strategy can limit carbon assimilation under prolonged stress, potentially reducing resilience. However, early stomatal closure may also be advantageous under severe or recurrent droughts, with drought conditions favoring different leaf strategies (Martínez-Vialta & Garcia-Forner 2017). In contrast, broad-leaved species generally maintain higher hydraulic efficiency, being able to sustain stomatal conductance for longer periods under water stress (Choat et al. 2012; Li et al. 2020). In this context, climate-driven shifts from needle-leaved dominated forests towards broad-leaved dominated forests have been already reported under the current climate change (Gregor et al. 2022; Ma et al. 2023). Therefore, despite the high interspecific variability, consistent biogeographical and functional patterns might emerge, offering valuable insights into forest responses to climate change.

Here, we seek to uncover broad patterns in forest species growth and mortality in response to climate change across large environmental gradients. To achieve this, we assessed the influence of climate and forest structure on growth and mortality for the 21 most widespread forest species in Europe across more than 50,000 forest inventory plots, covering from Mediterranean to boreal climates. We quantified the relative importance of climatic and structural drivers for each productivity component (growth and mortality occurrence and intensity) across species and functional groups, including broad-leaved and needle-leaved species. We then examined patterns of change in these components in response to spatial variation of vapour pressure deficit (VPD) anomaly, considering arid and wet species’ distribution edges as well as early and late stand developmental stages, aggregating responses across species and functional groups. We hypothesise that increased VPD anomalies will reduce tree species growth and increase mortality due to the combined effect of warming and water deficit stress (McDowell et al. 2020; Evans et al. 2025). We expect these effects to be stronger at the arid than the wet edges of the species’ distributions (Anderegg et al. 2019), and at early rather than late stand development stages (van Breugel et al. 2012; Máliš et al. 2023). Furthermore, we expect the observed patterns to reveal heightened sensitivity of needle-leaved species (Anderegg et al. 2020). This study aims to elucidate how the interaction between climate change, stand development and species’ climatic distribution affects forest demographic responses, potentially compromising the carbon sink function of European forests. We also aim to evaluate the importance of functional group- and species-level differences in shaping these responses, which is key for anticipating the effects of future changes in climate.

## 2. Methods

### 2.1. Forest Inventory data

We used National Forest Inventory data from five European countries covering Europe’s latitudinal gradient, selecting the most abundant tree species for analysis (see Appendix S1). National Forest Inventory data consists of standardised, plot-based repeated measurements recording tree species identity, size, and vitality status (alive or dead) for all trees within sampling plots, where plot radius may vary with tree size depending on the country (see Appendix S1 for additional country-level information). For our analyses, non-native species were excluded as well as species present in less than 1,000 plots across all countries. When a species was present in fewer than 50 plots within a specific country, that country was excluded from the analyses for that species to avoid issues associated with country-level low sample size. Although plot design, measurement characteristics and timeframe differed between countries (Appendix S1), all datasets consisted of two consecutive tree-level measurements, allowing the calculation of biomass gains and losses between measurements. The time interval between consecutive measurements ranged from 5 to 13.5 years depending on the country with an average of 7.5 years (Appendix S1). The maximum period covered considering all plots in our dataset was 22 years (1997-2019), with 2007 being the mean date for the first measurement and 2015 the mean date for the second measurement across countries (Appendix S1). We excluded plots with no adult trees alive in the first measurement and with unidentified tree species (1107 plots; 1.2%). We also excluded plots with records of harvest between measurements to avoid management-induced mortality from obscuring natural mortality responses (23,303 plots; 29%). We considered a plot as harvested when at least one tree was recorded as dead and absent in the second measurement. In the end, we analysed 52,703 plots and 706,352 adult trees (i.e. > 10 cm in diameter at breast height, d.b.h.) of the 21 most abundant tree species.

### 2.2. Productivity components: growth and mortality occurrence and intensity

We used consecutive forest inventory measurements to calculate annual productivity for the 21 forest species in our database. In each plot containing a target species, we calculated the target species stand-level growth and mortality as the main components of productivity (see Figure 1 for mapped observed values and Appendix S1 for species mean values).

**Figure 1.**
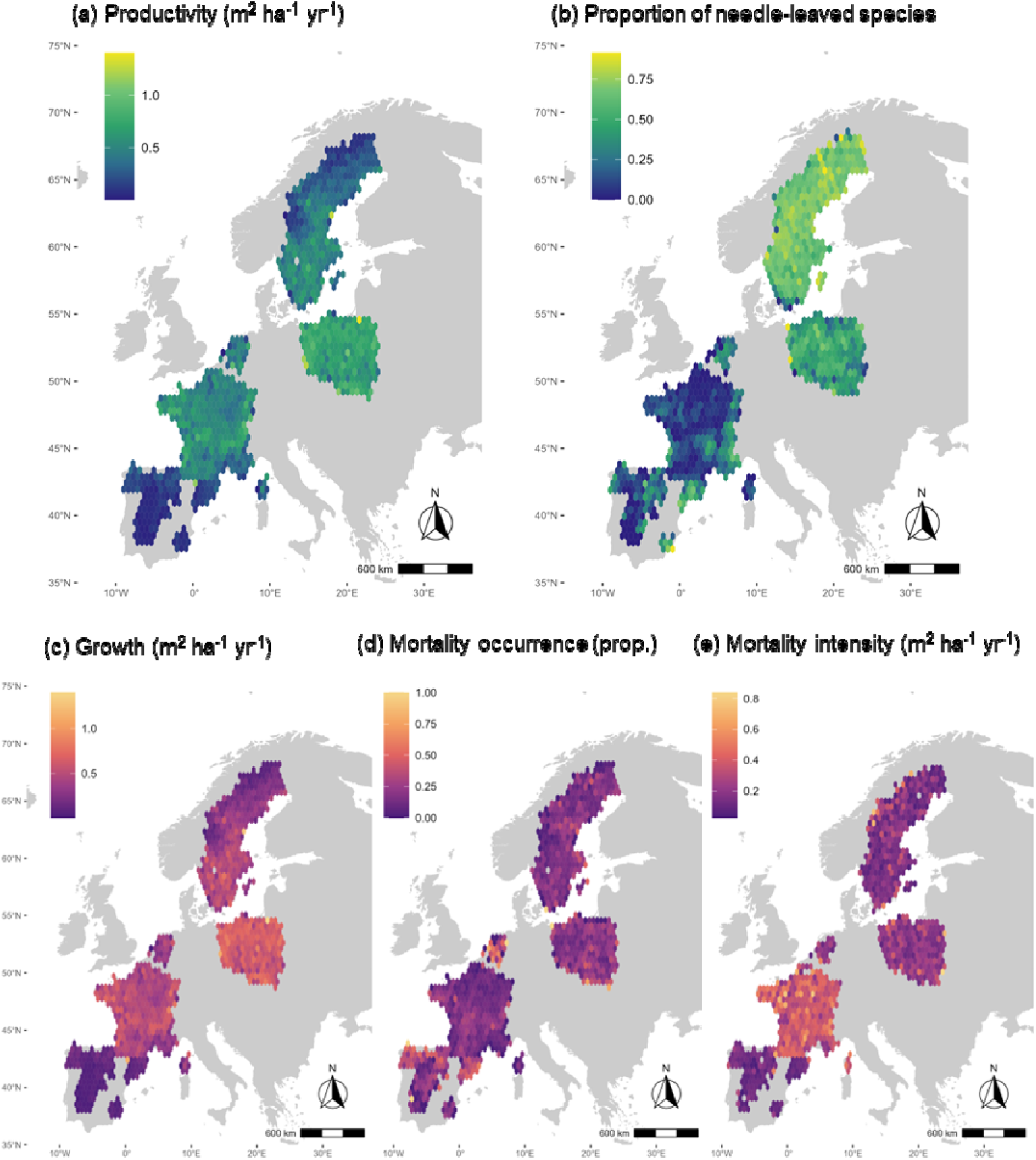
Spatial patterns of observed (a) median net forest productivity (i.e., growth minus mortality intensity), (b) proportion of needle-leaved species, (c) median forest growth, (d) proportion of mortality occurrence, and (e) median mortality intensity for all 21 target species per plot, averaged into ∼0.9° × 0.9° latitude–longitude grid cells to enhance visualization. Values above the 95^th^ percentile were excluded for all variables except mortality occurrence, since it is a binomial variable. See the distribution of structural and climatic variables in Appendix S2.

Species-specific stand-level growth (m^2^ ha^-1^ yr^-1^) was calculated as the sum of basal area increments of adult trees of the target species alive in consecutive inventories (i.e. adult growth) and the sum of the basal area of adult trees of the target species recruited in the second inventory (i.e. ingrowth), divided by the number of years between measurements. We included ingrowth as part of stand-level growth because it captures basal area gains associated with new adult trees, thereby contributing to the net productivity of the stand (Kloeppel et al. 2007). Species-specific stand level mortality was calculated as the sum of basal area of adult trees of the target species alive in the first inventory measurement but dead in the second. This variable exhibited a zero-inflated distribution, because most plots recorded no mortality between measurements (∼85% of the plots). Therefore, we distinguished between mortality occurrence and intensity. Mortality occurrence was calculated as a binomial variable where 0 represented no recorded mortality for the target species between measurements and 1 represented that at least one individual was recorded as alive in the first measurement and dead in the second in each plot. Mortality intensity (m^2^ ha^-1^ yr^-1^) was calculated for the plots in which mortality occurred for the target species and consisted of the basal area lost due to natural mortality (i.e. trees alive in the first inventory and dead but present in the second), divided by the number of years between inventories. Basal area was chosen as the reference population size metric to calculate forest growth and mortality intensity because it is directly calculated from tree size and is a widely available and precise measure across countries (Gschwantner et al. 2022).

### 2.3. Forest structure as predictor for forest growth and mortality

To characterise the most relevant drivers for tree species productivity related to stand structure, we used the first forest inventory measurement. From an initial selection based on the literature, we selected the most integrative and less correlated forest structural variables (see Appendix S2 for the complete initial set of variables calculated and further details on variable correlations). To account for the target species condition and intra-specific competition that strongly determines growth and mortality (Gómez-Aparicio et al. 2011; Idoate-Lacasia et al. 2025), we included target species abundance (i.e. target species basal area as the sum of the basal areas of all individuals of the target species in the plot, m^2^ ha^-1^). To consider the combined influence of intra- and interspecific competition on the components of forest productivity, we selected stand density (i.e. total number of trees per plot, No. trees ha^-1^), which has been shown to be an important predictor for forest dynamics (Ruiz-Benito et al. 2013; Zhang et al. 2017; Changenet et al. 2021). As a proxy for stand developmental stage (Ducey & Kershaw 2023), we calculated in each plot the mean diameter of the thickest 100 trees per hectare (mm ha^-1^).

Although we only selected plots without recorded harvesting during the study period, plots may have been influenced by past harvesting or harvesting in surrounding areas. To account for this, we included the harvest regime (% of the total tree basal area in the grid cell that was harvested annually) into our models as an integrative measure of harvest intensity and frequency at 1-degree grid resolution (Suvanto et al. 2025).

### 2.4. Aridity and recent climate change as predictors of forest growth and mortality

To characterise the climatic conditions in each plot, we used monthly climate data from CHELSA v2.1 (Karger et al. 2021) spanning from 1980 to 2018, a reliable climate data source for large spatial scale analyses (Rodríguez-Rey & Jiménez-Valverde 2024). We downloaded a set of water and temperature related variables and selected two low correlated variables known to drive species productivity dynamics: Climate Moisture Index (CMI, kg m^-2^) and Vapour Pressure Deficit (VPD, Pa). CMI is the difference between precipitation and potential evapotranspiration and serves as a robust indicator of aridity, driving forest growth and mortality (Berner et al. 2017; Hogg et al. 2017; D’Orangeville et al. 2018; Luo et al. 2019). VPD is the difference between the saturation vapor pressure and actual vapor pressure (Grossiord et al. 2020; Lansu et al. 2020), and it is an increasingly used climate change metric that drives productivity and forest mortality in both high and low latitudes (Castellaneta et al. 2022; Hammond et al. 2022; McDowell et al. 2022).

To characterise the reference aridity gradient across Europe, we calculated the mean CMI for each plot over a 15-year reference period from 1980 to 1995. This period was chosen to avoid overlap with our study period, which started in 1997 for some plots. For each year of the reference period, we calculated the annual mean CMI from monthly data and then averaged these annual values to obtain a single mean CMI value per plot for the reference period. Lower values of CMI indicate greater aridity.

To evaluate the recent climate change experienced by each plot during the study period (i.e. years between the first and second forest inventory measurements) relative to the reference period (1980-1995), we calculated a standardised VPD anomaly (unitless) focusing only on the growing season months (March-August). Using a uniform growing season (March–August) across all plots, allowed us to maintain consistency in the calculation of climate anomalies while capturing the primary period of forest growth and atmospheric water demand across Europe. We calculated VPD anomaly using the same methodology as in the Standardised Anomaly Index first developed by Kraus (1977) as follows:

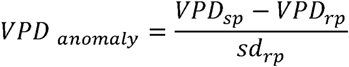

where *VPD_sp_* is the mean VPD of the study period of each plot, *VPD_rp_* is the mean VPD of the reference period and *sd_rp_* is the standard deviation of *VPD_rp_*. Since the study period differed between plots (i.e. measurement years differed between countries and not all plots within the same country were measured in the same year), *VPD_sp_* was calculated specifically for the study period of each plot. Higher values of VPD anomaly indicate an increase in VPD for the study period relative to the reference period, reflecting increased transpiration pressure due to climate change.

### 2.5. Species-specific forest growth and forest mortality models

For each of the 21 target species and productivity components (i.e. growth and mortality occurrence and intensity) we fitted generalised additive models (GAM) using *mgcv* R package (Wood 2011) in R version 4.4.2 (R Core Team 2024). This approach allowed us to evaluate species-specific productivity responses to stand development and aridity under recent climate change.

We fitted GAM specifying a Gamma distribution with a log link function to model the continuous right-skewed distribution of forest species growth and mortality intensity. Mortality occurrence was modelled as a binary response using a binomial distribution with a logit link function. Target species basal area, stand density and harvest regime were included as linear predictors in the parametric component of the models based on their ecological relevance. Country was also included as a fixed categorical effect inside the parametric component of the models to account for systematic differences in measurement protocols across national inventories. Species basal area and stand density were log-transformed to linearise the relationship with the response variables. The interactive effects of stand development, aridity and VPD anomaly were modelled in the non-parametric component using a three-way tensor product smooth with thin plate regression splines (*bs = “tp”*). The thin plate regression splines provide high stability and flexibility for multidimensional smooths and is recommended when interacting variables differ in magnitude (Wood 2017). The triple interaction was modelled using a non-parametric smooth because we hypothesized that the combined effect of these three variables on the response would be complex and non-linear, and quantifying this non-linearity was a central objective of the study. In contrast, other covariates were included as parametric terms after appropriate transformations to achieve approximate linearity, as their primary role was to control for confounding effects rather than to explore potentially non-linear relationships. Pairwise Pearson correlations among predictors were weak (|r| < 0.3, see Appendix S2), indicating limited linear dependence. Visual inspection of the joint three-way covariate space showed no major gaps across the observed predictor range (Appendix S2), supporting the implementation of higher-order smooth interaction terms.

To assess the models’ fit, we simulated the residuals and evaluated diagnostics plots with *DHARMa* (Hartig & Hartig 2017) and *gratia* (Simpson 2024) R packages. To report the goodness of fit, we extracted the explained deviance of each model (Lai et al. 2024). Predictor importance was evaluated using the change in deviance, which quantifies the increase in deviance when a given predictor is removed from the model, following the leave-one-covariate-out (LOCO) approach (Vakhitova & Alston-Knox 2018; Hayes 2021; Bladen et al. 2024). Besides assessing model fit, predictive performance was also evaluated as an additional validation. For this purpose, we applied cross-validation using 500 random subsampling iterations. In each iteration, 80% of data was randomly selected for training and the remaining 20% for testing (Gelman et al. 2021), consistent with standard cross-validation protocols (Kuhn & Johnson 2013). Our approach provided an assessment of model stability and predictive inference, with model results subsequently weighted according to model predictive performance.

For forest growth and mortality intensity models, Root Mean Square Error (RMSE) and Mean Absolute Error (MAE; Willmott & Matsuura 2005) were used as predictive performance metrics computed using *yardstick* (Kuhn et al. 2025) R package. For comparability among species, normalised RMSE (NRMSE) and Normalised MAE (NMAE) were calculated by dividing each value by the range of the response variable in the model. Both RMSE and MAE provide information on model fit and predictive accuracy (Li 2016), being MAE less sensitive to outliers than RMSE, but both indicating greater predictive performance if their values are small (Chicco et al. 2021). For forest mortality occurrence models, we evaluated model performance using the area under the receiver operating curve [AUC, (Fielding & Bell 1997)] and Se* (Jiménez-Valverde 2014). AUC is a widely used performance metric whereas Se* is a novel discrimination metric which accounts for some of the AUC limitations related to prevalence dependence (Jiménez-Valverde 2014, 2020). A combination of metrics was used to minimise metric-specific bias using the *ROCR* (Sing et al. 2005) and *caret* (Kuhn 2008) R packages. AUC is a threshold-independent metric ranging from 0 to 1, with values below 0.5 indicating no discriminative ability while values of 1 represent perfect classification. Se* represents the overall probability of correctly classifying any given case and uses the threshold that minimises the difference between sensitivity and specificity, with greater values of Se* indicating a better classification accuracy.

### 2.6. Species forest growth and mortality predictions along VPD anomaly

To predict changes in species growth and mortality along the VPD anomaly gradient, we generated model-based predictions from each species-specific GAM while holding stand development and aridity constant at fixed levels. Specifically, for each species we specified two levels for stand development (i.e. early stand development as the 25^th^ percentile and late stand development as the 75^th^ percentile observed for each species in our database), two levels for aridity (i.e. arid using the 15^th^ percentile and wet using the 85^th^ percentile of the CMI observed within each species full distribution in Caudullo et al. (2017)) and a set of fixed VPD anomaly values defined at regular intervals of 0.001 over the observed VPD anomaly range for each species (i.e. from the minimum to the maximum value observed in our dataset for each species). By estimating aridity using each species’ full European distribution rather than only the subset of their distribution represented in our database, the models’ predictions more accurately reflected the climatic conditions experienced across their entire biogeographical ranges. To gain further insight into the relationship between aridity and VPD anomaly, we also examined the effect of VPD anomaly under mild aridity conditions (i.e. using the 50^th^ percentile of the CMI observed within each species full distribution in Caudullo et al. (2017)). Predictions were obtained for the full factorial combination of the covariate levels. For additional explanations of the definitions of aridity levels, see Appendix S4.

Species-level predictions were aggregated to obtain overall mean responses and functional group–specific responses (broad-leaved vs. needle-leaved) for each productivity component. Aggregation across species was performed using a weighted mean based on the models’ predictive performance (i.e., MAE for continuous responses or AUC binary responses). Thus, species with lower predictive accuracy contributed less to the final estimate. We calculated these weights using a SoftMax function with temperature parameter (τ), a method commonly used to convert performance metrics into importance weights while controlling the sharpness of the weighting distribution with the τ value (Totaro et al. 2020; Pretis & Landes 2021; Khan et al. 2023). We set τ = 0.4 to ensure relatively uniform weights and to avoid excessive differences among species and prevent excessive dominance of any single species in the aggregated predictions (see Figure S1 for a sensitivity analysis using τ values of 0.2, 0.6 and 1).

We also assessed species-specific growth and mortality responses along VPD anomaly extremes by predicting species-level growth and mortality (occurrence and intensity) at the minimum and maximum VPD anomaly values observed for each species. Then, we quantified species-specific sensitivity to VPD anomaly as the difference between predictions at the maximum and minimum VPD anomaly values at each species’ aridity edge and stand development stage. Positive values indicate an increase in the target response (i.e. tree growth, mortality occurrence or intensity) with higher VPD anomaly, whereas negative values indicate a decrease. To assess whether species sensitivity was statistically significant, we calculated the 95% confidence interval of the difference between predictions at maximum and minimum VPD anomaly (±1.96 standard errors of the predictions). Species were considered to exhibit a significant increasing or decreasing response when this confidence interval did not overlap zero (Gelman et al. 2021).

## 3. Results

### 3.1. Relative importance of forest structure, climate and management for species’ aboveground productivity

Forest growth and mortality drivers showed a high interspecific variability in their relative contribution to the explained deviance. Variability was higher for mortality than growth, especially for mortality intensity (Figure 2a). Despite the high variability, some common patterns were found for forest growth and mortality between individual species (Figure 2a), as well as between broad-leaved and needle-leaved species (Figure 2b). Forest structure was overall the most important factor influencing both growth and mortality, although the relative importance of its components varied between them. The target species basal area was most influential in mortality occurrence models, whereas both target species basal area and stand development were important for forest growth and mortality intensity (Figure 2a). Stand density was relatively more important for mortality than growth (Figure 2a). Climate played a less dominant role than forest structure, but it was more relevant for mortality than for growth, particularly for mortality intensity (explained deviance ∼38% and ∼15% for mortality intensity and occurrence respectively in contrast to ∼5% for growth in average; Figure 2a); notably, for five species, climate was the main driver of forest mortality intensity (Figure 2a). These patterns were consistent across functional groups, with climate playing a relatively larger role in needle-leaved than broad-leaved species (Figure 2b).

**Figure 2.**
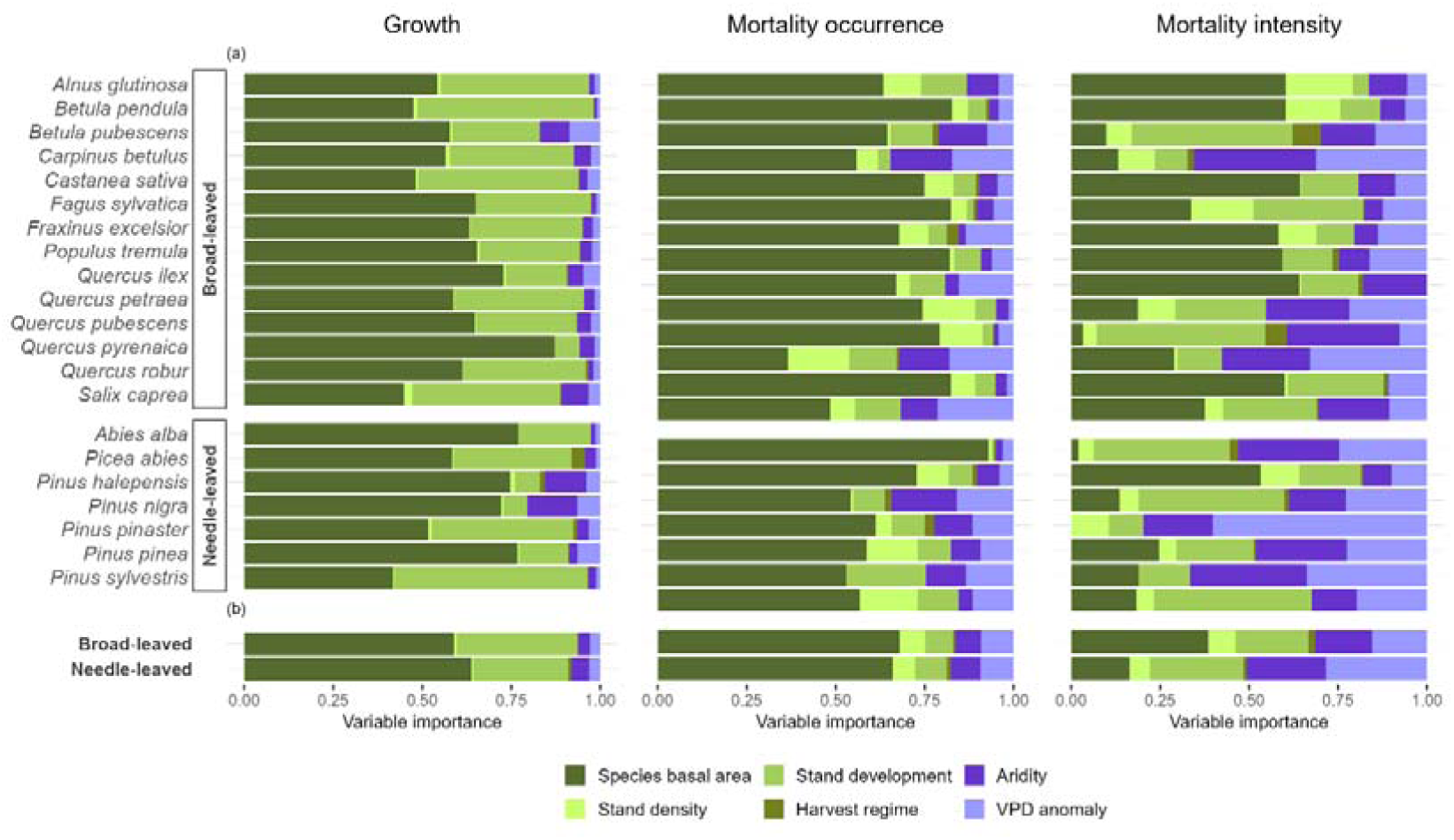
(a) Proportion of deviance explained by forest structure (i.e. species basal area, stand development and stand density), climate (i.e. aridity and VPD anomaly) and harvest regime (i.e. intensity and frequency of harvest) for the species growth, mortality occurrence and mortality intensity models, calculated as the explained deviance lost when each predictor was removed from each species’ model. (b) Mean explained deviance for broad-leaved and needle-leaved species weighted by the predictive accuracy of each species model (NMAE for forest growth and mortality intensity models and AUC for the mortality occurrence model).

Diagnostics plots indicated overall good fit of forest growth and mortality models (Appendix S3), with residuals displaying no major deviations from model assumptions. The explanatory power varied substantially among species and productivity components (Appendix S3). The highest explained deviance was observed for growth models (0.219-0.682; mean 0.483), intermediate for mortality intensity models (0.187-0.595; mean 0.339) and the lowest for mortality occurrence models (0.135-0.318; mean 0.203). Furthermore, the predictive accuracy differed among species, with growth models showing acceptable performance, whereas mortality intensity models exhibited greater uncertainty (i.e. most NMAE growth values < 0.5 and mortality intensity < 0.7; Appendix S3). Mortality occurrence models showed average to acceptable predictive accuracy, with all species exhibiting mean AUC values above 0.75 and mean Se* values exceeding 0.68 (Appendix S3).

### 3.2. Predicted changes in species productivity components along gradients of VPD anomaly, aridity and stand development

The three-way interaction between VPD anomaly, stand development and aridity strongly influenced species-level growth and mortality occurrence and intensity (Figure 3–4; see Appendix S5 for species-specific responses). When quantifying sensitivity as the difference between predicted responses at the minimum and maximum species VPD anomaly values, only a subset of species showed statistically significant differences based on the 95% confidence interval (±1.96 standard errors; Figure 3). The overall directional trends shown in Figure 3, although weak, showed declining growth and increasing mortality occurrence and intensity with increasing VPD. Specifically, most growth responses were negative (i.e. bars and numbers located to the left of the dashed line), indicating reduced growth under the highest observed VPD anomaly relative to the lowest observed VPD anomaly, whereas most mortality responses were positive (i.e. bars to the right of the dashed line), indicating increased mortality under the highest observed VPD anomaly relative to the lowest observed VPD anomaly. These patterns become clearer when applying a less restrictive confidence interval (±1 standard error; Appendix S4).

**Figure 3.**
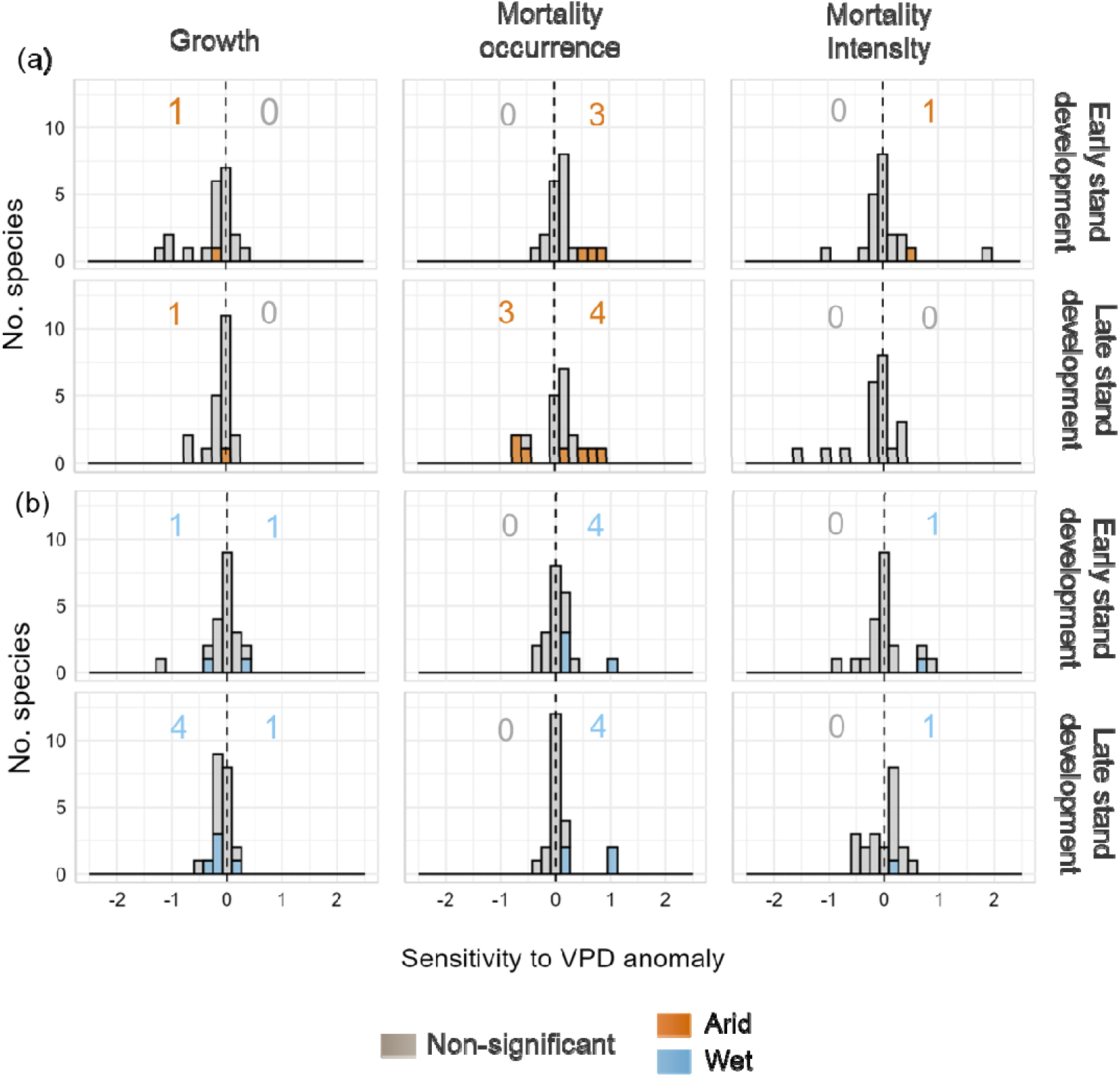
Species-specific sensitivity to increased VPD anomaly in (a) arid (orange) and (b) wet (blue) edges of species’ distributions and across early and late stand developmental stages. Sensitivity to VPD anomaly was calculated for each species as the difference in their response between its maximum and minimum VPD anomaly values, with bars and numbers to the right of the dashed line showing an increase in the species response with higher VPD anomaly and those to the left showing a decreased response. Coloured bars and numbers represent species for which climatic sensitivity was statistically significant, with the significance being determined by whether the 95% confidence interval of the difference in the response between high vs. low VPD anomaly (±1.96 standard errors) at the species level excludes cero.

**Figure 4.**
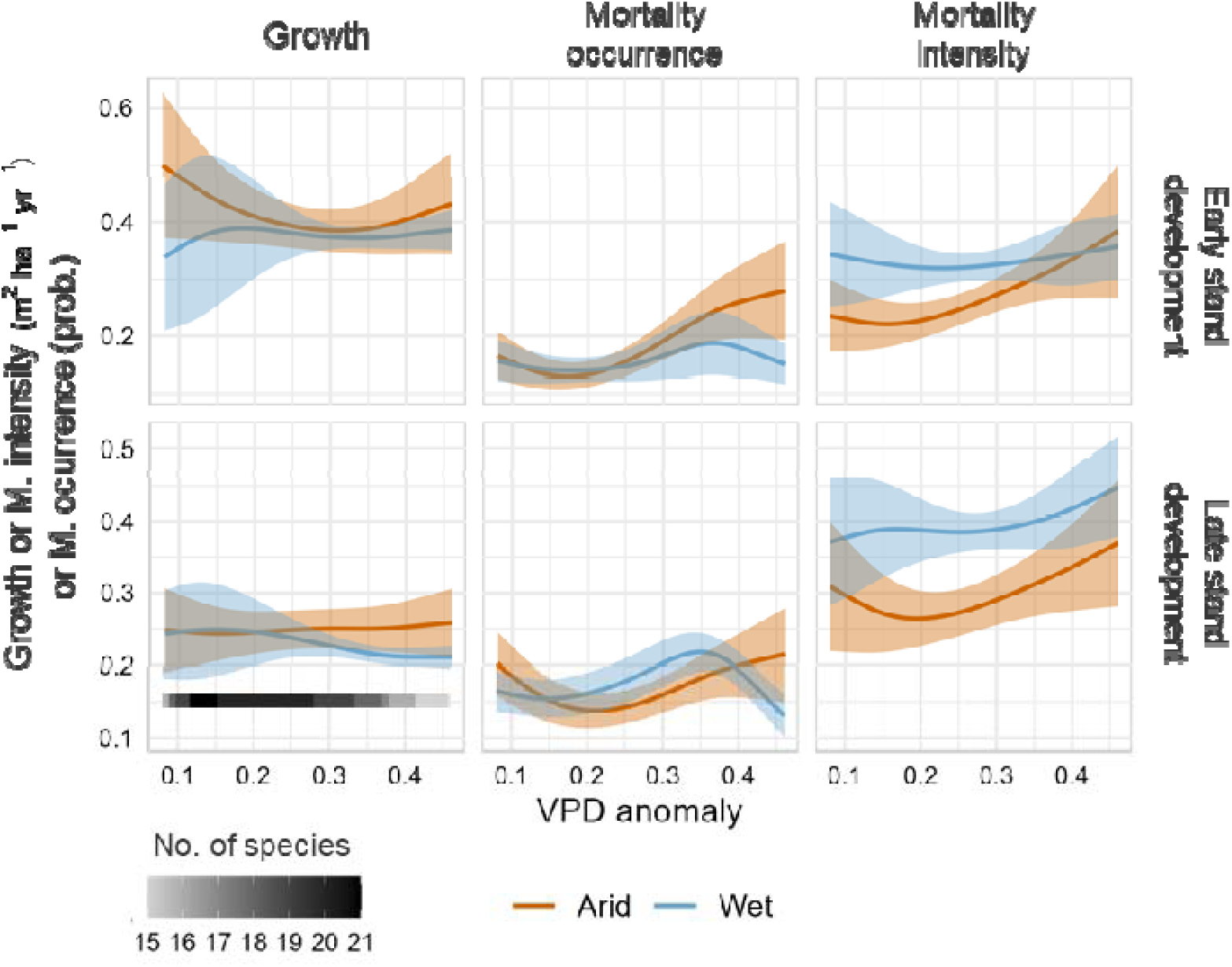
Weighted mean species models predictions for productivity components along the VPD anomaly gradient, restricted to the range where at least 75% of all species were present. Each species’ contribution to the mean prediction was weighted by the predictive accuracy of its model, NMAE for growth and mortality intensity models and AUC for mortality occurrence models. Shaded areas represent 95% confidence intervals derived from the weighted standard error of the species-specific predictions. The number of species contributing to the mean prediction within each segment of the VPD anomaly gradient is indicated by shading from white to black in the legend. Predictions were computed at the 25^th^ (i.e. early) and 75^th^ (i.e. late) percentiles of stand development for each species, and the species-specific arid or wet edge at the 15^th^ (i.e. arid in orange) and the 85^th^ (i.e. wet in blue) percentiles of climate moisture index along the observed species’ VPD anomaly range.

When examining growth and mortality patterns across the full VPD anomaly gradient, rather than focusing only on the extremes, non-linear patterns emerged (Figure 4). In addition, aggregated species-level growth and mortality responses showed contrasting patterns between the arid and wet edges of species’ distributions and between early and late stand developmental stages (Figure 4). In particular, the predicted changes in forest growth and mortality occurrence and intensity along the VPD anomaly gradient were generally more pronounced at the arid edge than at the wet edge of the species’ aridity distribution, particularly for early stand developmental stages (Figure 4). This pattern remained when analyses were restricted to models including only significant three-way interactions among aridity, stand development and VPD anomaly (Figure S2). Under mild aridity conditions within species’ distributions, growth and mortality responses to increasing VPD anomaly followed patterns similar to those observed at the arid and wet distributional edges, with mortality increasing and growth declining as VPD anomaly increases (Appendix S4).

### 3.3. Predicted changes in productivity components for broad-leaved and needle-leaved species

Forest growth and mortality responses to VPD anomaly were non-linear and generally exhibited higher uncertainty and interspecific variability in needle-leaved than in broad-leaved species (see higher variability among needle-leaved species in Appendix S4, Figure 5), leading to generally weak aggregated patterns for needle-leaved species growth and mortality intensity responses, but not for mortality occurrence with increasing VPD anomaly (Figure 5). The most pronounced responses to VPD anomaly for broad-leaved species occurred in arid edges and early stand development forests for the three productivity components (Figure 5a), while for needle-leaved they occurred in wet edges for mortality occurrence (Figure 5b). Furthermore, the highest differences between needle-leaved and broad-leaved species occurred for forest mortality in wet edges (Figure 5b).

**Figure 5.**
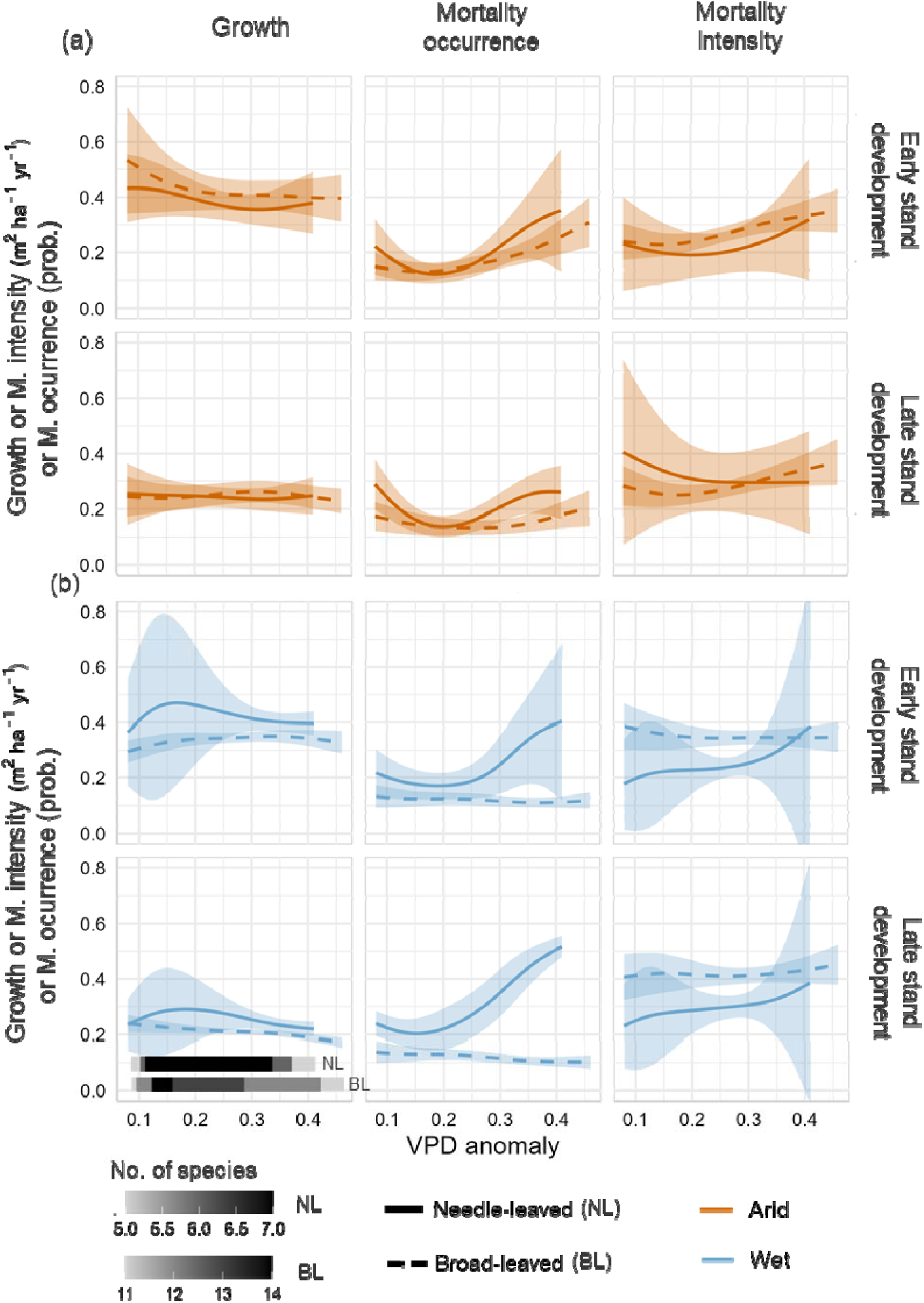
Weighted mean broad-leaved and needle-leaved species predictions of productivity components by functional group along the VPD anomaly gradient, restricted to the range where at least 75% of species were present. Each species’ contribution to the mean contribution was weighted by the predictive accuracy of its model, NMAE for growth and mortality intensity models and AUC for mortality occurrence models. Shaded areas represent 95% confidence intervals derived from the weighted standard error of the predictions. The number of species contributing to the mean prediction within each segment of the VPD anomaly gradient is indicated by shading from white to black in the legend. Predictions were computed at the 25^th^ (i.e. early) and 75^th^ (i.e. late) percentiles of stand development for each species, and the species-specific’ arid or wet edge at the 15^th^ (i.e. arid in orange) and the 85^th^ (i.e. wet in blue) percentiles of climate moisture index along the observed species’ VPD anomaly range.

## 4. Discussion

Using a comprehensive database of National Forest Inventories across Europe, we observed high variability in the relative importance of forest structure and climatic drivers of forest growth and mortality across the 21 most abundant tree species. However, some common patterns emerged across species, highlighting forest structure-driven responses in most species, with a greater importance of climate in mortality (especially mortality intensity compared to occurrence), and needle-leaved species compared to broad-leaved species. The pronounced interspecific differences in growth and mortality responses to increasing climatic stress suggest that robust future projections of forest dynamics will require explicit consideration of separate productivity components (Benito-Garzón et al. 2013; Astigarraga et al. 2020; McDowell et al. 2020; Kunstler et al. 2021) and species-specific assessments (Ruiz-Benito et al. 2013; Changenet et al. 2021; Astigarraga et al. 2024). Furthermore, we evidenced forest growth declines and forest mortality increases along VPD anomaly gradients at large spatial extents, particularly in early stages of development at arid edges of the species’ distributions. Our results further indicate that at arid edges of the species’ distributions, populations more pronounced responses to climate change than at wet edges, especially for needle-leaved species and early stand development forests. The heightened susceptibility to increased VPD anomaly observed at arid edges underscores the importance of temperature-driven stress in shaping vegetation dynamics (Evans et al. 2025).

### 4.1. Climate as a key underlying driver of mortality, despite high variability among species and forest structure importance

Forest structure emerged as the most relevant driver of species growth and mortality (Figure 2). The generally highest importance of forest structure aligns with the major role of stand biomass and forest structure on forest dynamics found by previous studies (Benito-Garzón et al. 2013; Zhang et al. 2015; McDowell et al. 2020; Zhai et al. 2024). However, we found a considerable variability in the contribution of the individual structural variables among species and forest productivity components (Figure 2). Stand density had a stronger influence on forest mortality than on growth, in agreement with the key role of competition in mortality processes (Van Gunst et al. 2016; Jump et al. 2017a; Kulha et al. 2023). Stand development was particularly relevant for growth and mortality intensity, but less so for mortality occurrence. Despite stand development being a key driver of forest dynamics (Ruiz-Benito et al. 2015; Astigarraga et al. 2024; Bordin et al. 2025), its effect was lower for mortality occurrence than intensity. It could be due to the fact that mortality occurrence could be more stochastic and related to background mortality (i.e., mortality in the absence of catastrophic events; Franklin 1987; Csilléry et al. 2013), whereas mortality intensity can be more related to climate-induced tree mortality (e.g. die-off events where many trees are damaged or death as a response to exceptional events, Hammond et al. 2022; Rebollo et al. 2024).

Climate had a higher importance on forest mortality than on growth (climate explaining a deviance c. 30% and 10% higher for mortality intensity and occurrence, respectively, than for growth; Figure 2). The greater importance of climate for mortality than for growth is consistent with previous findings showing that forest mortality responses to climatic stress can be abrupt due to the surpassing of physiological thresholds, under the interaction of multiple factors such as drought, warming, insect outbreaks and forest structural conditions (Anderegg et al. 2015; Hammond et al. 2022). Furthermore, the low deviance explained by climate for forest growth may reflect the particularly strong influence of forest structure, as well as contrasting effects of climate change on growth, including both negative and positive effects associated with, for example, elevated CO_₂_ (Kauppi et al. 2014; Grossiord et al. 2022; Zhong et al. 2023). In addition, the strong effect of climate on forest mortality might be underlined by the heterogeneity and stochasticity of both climatic events and mortality processes (Franklin et al. 1987; Benito-Garzón et al. 2013), which likely contributes to the high explained deviance of climate on forest mortality and the lowest explanatory and predictive power of mortality models compared to growth models.

Climate was the most relevant predictor of mortality intensity for five of the 21 species, but it also exhibited the highest species-level variability in driver importance. The high species variability might reflect differences in climate among species, and a strong dependence on species-specific strategies and evolutionary history (Barrere et al. 2023), as it is partially observed in the higher importance for needle-leaved vs. broad-leaved species (Figure 5). The stronger influence of climate on mortality intensity than on occurrence aligns with results of previous studies suggesting than the former is more related to die-off events and the latter to background forest mortality (Das 2016; Changenet et al. 2021). In addition, temperature-related variables as VPD anomaly may capture the effects of extreme events, such as heatwaves or severe droughts due to compound events (e.g. warming increasing the probability of hotter droughts, Bastos et al. 2021). Compound events are also associated with high-intensity mortality rather than with background mortality (Gazol & Camarero 2022). Together, these results highlight the key role of climate-driven mortality in shaping long-term forest dynamics (McDowell et al. 2020). They also emphasise the importance of explicitly accounting for both mortality occurrence and intensity when assessing forest productivity responses, as they may be influenced differently by specific drivers and contribute unequally to changes in forest productivity (Changenet et al. 2021), particularly given the disproportionately large impact that mortality can have on forest productivity (Anderegg et al. 2016; Astigarraga et al. 2020).

### 4.2. Decreased growth and increased mortality along VPD anomaly gradients were strongest at arid species’ edges and early developmental stages

Forest growth declined and mortality increased along the VPD anomaly gradient as hypothesised (Figure 4), with most of the species following the expected pattern in both aridity edges and stand developmental stages. However, only few tree species exhibited significant differences between their VPD anomaly extremes (see species-specific responses in Figure 3 and in Appendix S5), which might be due to forest growth and mortality responses not being linear along increasing VPD anomaly, as evidenced in Figure 4 and when looking at species-specific trends (Appendix S5). The effect of VPD anomaly on the productivity components might be likely driven by the rapid rise in VPD in the last decades, which in fact is outpacing the rate of increase in CO_₂_ and temperature (Song et al. 2022). Increased VPD is strongly linked to plant stress because it directly reflects atmospheric and soil humidity, thereby impacting forest growth and mortality through carbon starvation and hydraulic failure (Yuan et al. 2019; Grossiord et al. 2020; McDowell et al. 2022). Additionally, the patterns of increasing forest mortality with increasing VPD could be due to compound climatic events, as warming has been shown to increase the probability of other extreme climatic events that are detrimental to forest productivity such as droughts (Reyer et al. 2017; Seidl et al. 2017; Hartmann et al. 2022). However, the observed increase in forest growth with rising VPD in wet edges and early stand development stages may reflect the positive effects of warming for species typical of temperature-limited forests (Kauppi et al. 2014; Pretzsch et al. 2014; Green et al. 2020), fertilisation effects and the lengthening of the growing season (Park et al. 2016; Gao et al. 2022; Song et al. 2022). The increased forest growth observed in wet and early stand-development forests align with previous evidence of increased productivity and growth sensitivity in young and recently stablished forests (Anderson-Teixeira et al. 2013; Alfaro-Sánchez et al. 2019). However, the fact that some species still exhibited the hypothetised decline in growth even at wet edges suggest that certain species might be more vulnerable to warming than others (Fei et al. 2017; Astigarraga et al. 2024). Although forest growth generally decreased and mortality increased along the VPD gradient across all edges and stand developmental stages, differences in the magnitude of these responses were found both at the species-level and in aggregated responses.

The most pronounced responses to VPD anomaly occurred at arid compared to wet edges of species’ distributions and in early stand-development forests (Figure 4). While the interaction between aridity, stand development and VPD anomaly was not significant for some species, the overall pattern was observed independently of including or excluding these tree species, supporting the robustness of the results (Figure S2). Our results are consistent with early successional stands experiencing higher competition and having weaker climatic buffering capacity (van Breugel et al. 2012; Máliš et al. 2023), which could lead to greater vulnerability to water limitations (Ruiz-Benito et al. 2014). The fact that forest growth and mortality responses to VPD anomaly were most pronounced at the arid edges of species distributions agrees with previous studies (Changenet et al. 2021; Astigarraga et al. 2024). This might suggest that peripheral populations at the rear edge are responding differently in terms of forest growth (Kunstler et al. 2021) and background mortality (Archambeau et al. 2020; Changenet et al. 2021). In contrast, die-off events are being observed along the entire species distributional range but show an increasing probability towards high VPD anomalies (Jump et al. 2017; Changenet et al. 2021). The fact that mortality intensity responses to rising VPD occur under many different climatic and structural conditions suggests that other factors, such as increased density or stand age, further modulate responses to changing climate (Jump et al. 2017).

### 4.3. Susceptibility to increased VPD anomaly was greater for needle-leaved than for broad-leaved species

Climate had a stronger effect on forest growth and mortality for needle-leaved than for broad-leaved species, explaining a higher amount of deviance and suggesting a higher sensitivity of needle-leaved species to increased VPD (Figure 2). The stronger effect of climate in needle-leaved than broad-leaved species is consistent with expected climate-driven differences between these two functional groups (Cailleret et al. 2017; Blackman et al. 2019; Zhao et al. 2024; Pompa-García et al. 2025). Several studies have suggested that needle-leaved species exhibit greater sensitivity to climate-related stress (Greenwood et al. 2017; Anderegg et al. 2020; Barrere et al. 2023), which has been linked to differences in leaf traits, xylem hydraulics and carbon-use strategies (Johnson et al. 2012; Li et al. 2020), as well as to contrasted strategies for coping with rising VPD (Johnson et al. 2012; Carnicer et al. 2013). On the one hand, needle-leaved species generally exhibit a conservative water-use strategy characterised by early stomatal closure to prevent hydraulic failure (McDowell et al. 2008). While this strategy may be effective under short-term stress, it might limit carbon assimilation, increasing vulnerability, under prolonged stress (Choat et al. 2012; Li et al. 2020). On the other hand, broad-leaved species tend to show more resistance to water limitation through higher cavitation resistance and hydraulic efficiency, sustaining stomatal conductance for longer periods under water scarcity and thereby maintaining growth and competitive ability under sustained stress (Choat et al. 2012; Li et al. 2020). In addition, the higher susceptibility of needle-leaved species could be amplified by land-use legacies, such as their frequent establishment in highly dense and low diverse plantations, which further increase their vulnerability to climate-driven stress (Ruiz-Benito et al. 2012; 2013; Lázaro-Lobo et al. 2022; Vilà-Cabrera et al. 2023). Our results also align with previous studies predicting climate driven shifts from needle-leaved dominated forests towards broad-leaved dominated forests under climate change (Gregor et al. 2022; Ma et al. 2023).

Needle-leaved species showed more non-linear responses with higher uncertainty along VPD anomaly gradients than broad-leaved species (Figure 5). The strongest responses to the VPD anomaly gradients were observed for needle-leaved species mortality occurrence, showing more pronounced responses, particularly at wet species’ edges and in late developmental stages, where the largest differences between functional groups were also detected. Furthermore, despite greater absolute growth and lower mortality intensity in the arid than wet edge, suggesting that many of these species might not be at their physiological limit, broad-leaved species exhibited the strongest responses to VPD anomaly at arid edges and in early developmental stages. The fact that broad-leaved species had most pronounced responses at their arid edge could suggest that despite potentially being less sensitive to climate than needle-leaved species, they could be more sensitive to increased temperature at their arid edge (Münchinger et al. 2023). However, the pronounced sensitivity of needle-leaved species mortality to VPD anomaly is consistent with their previously noted high susceptibility to climate, and aligns with studies documenting more pronounced responses to water stress and higher mortality rates in needle-leaved than in broad-leaved species (Greenwood et al. 2017; Ruiz-Benito et al. 2017b; Sáenz-Romero et al. 2019). The contrasting responses between functional groups likely reflect species-level differences. For example, our results suggest that some Mediterranean needle-leaved species might not be strongly constrained at their arid edge. In particular, *Pinus pinea* and *Pinus pinaster* showed declining mortality occurrence with increasing VPD anomaly at arid edges in late stand development stages, but high mortality occurrence at wet edges (Appendix S5). These patterns are consistent with previous findings reporting decreased mortality with increasing temperature for these species (Ruiz-Benito et al. 2013), which may reflect their greater adaptation to climatic stress than other temperate species, as previously observed in their demographic responses to climate (Benito-Garzón et al. 2013). Furthermore, a strong effect of climate change has been reported for needle-leaved species in the core of their distribution, potentially linked to legacy effects of high-intensity management (Jump et al. 2017).

### 4.4. Implications for forest dynamics under climate change

While productivity gains generally exceeded losses across the five European countries, implying continued carbon sink function at large spatial extents, our results indicate a strong climatic influence on mortality intensity (Changenet et al. 2021), and reveal reduced growth and increased mortality along VPD anomaly gradients. Our findings suggest that, if current climate change trends persist, climate-driven increases in tree mortality might become a major driver of large-scale biomass losses. Importantly, our dataset extends only to 2018, preceding a period of substantial VPD increases as a result of extreme drought events (Novick et al. 2024), suggesting that current and future short.-term responses could be even more pronounced. Moreover, the strongest sensitivities to increasing VPD anomaly were detected at the arid margins of species’ distributions, indicating that these populations may be particularly vulnerable to ongoing climate change. Conservation and adaptation strategies should therefore prioritise monitoring and assessment of forest responses in these areas of the species’ distributions. In addition, early stand development stages had heightened vulnerability to increases in VPD anomalies, highlighting the importance of protecting and maintaining mature forest, which can provide high buffering capacity against climate change (van Breugel et al. 2012; Máliš et al. 2023). This is particularly challenging in Europe, where old-growth forests are scarce due to historical land-use changes and intense forest management.

Our results highlight the need for species- and demographic-process-specific modelling to accurately assess climate change impacts on forests and improve projections of future forest dynamics. The large interspecific variability observed both across and within productivity components suggests that demographic responses cannot be fully captured using functional-grouped averages or mean-based approaches alone (Fei et al. 2017; Idoate-Lacasia et al. 2025). Furthermore, the non-linear nature of growth and mortality responses to VPD anomaly highlights the need for caution when assuming linear relationships between demographic processes and climate change. Species not only differed in the magnitude of their responses, but also in their shape and direction of responses along the climatic gradient, reinforcing the importance of explicitly accounting for species-specific responses. Overall, while broad-scale assessments of forest productivity provide important insights (Hogan et al. 2024), we evidence that robust projections of forest dynamics might require large-scale and species-level demographic data that separately quantify growth and mortality processes (International Forest Mortality Network 2025). In this context, our results provide benchmark information on key forest productivity components for the 21 most common tree species across Europe that could be used to further develop, evaluate and validate process-based demographic models of forest responses to ongoing climate change.

## 5. Acknowledgements

This research was funded by European Research Executive Agency (REA) through the Horizon Europe CLIMB-FOREST Project (No. 101059888). Views and opinions expressed are however those of the author only and do not necessarily reflect those of the European Union. The European Union cannot be held responsible for them. We also acknowledge support from are funded by the Science and Innovation Ministry, AEI (Agencia Estatal de Investigation) (subproject LARGE, N° PID2021-123675OB-C41). M.B.-H. was supported by a FPU fellowship of the Spanish Ministerio de Ciencia, Innovación y Universidades (MICIU) by the reference FPU23/00814. J.A. was supported by the Basque Government’s Postdoctoral Programme for the Improvement of Doctoral Research Staff (POS_2024_1_0026). A.V.-C. was supported by the Ministry of Science, Innovation and Universities (Spain) – Ramón y Cajal fellowship (RYC2023-045604-I). V.C.A. was supported by the Community of Madrid under the 2024 call for the ‘César Nombela’ research talent attraction programme (2024-T1/ECO-31335). C.G.-A was supported by funding from the Talent Attraction Programme of the Community of Madrid (2022-T1/AMB-23790). We thank all the organizations and NFIs who provided the data for this study. We thank the Ministerio para la Transición Ecológica y el Reto Demográfico (MITECO) for open access to the Spanish Forest Inventory (https://www.miteco.gob.es/), the Institut national de l’information géographique et forestière (IGN) for open access to the French Forest Inventory (https://www.ign.fr/) and the Wageningen University & Research for open access to the Dutch National Forest Inventory (https://www.wur.nl/). The National Forest Inventory in Poland had been funded by the State Forests Holding (“Lasy Państwowe”).

## 6. Authors contributions

M.B.-H, J.A., A.V.-C., T.A.M.P., and P.R.B conceived the study. M.B.-H, J.A., C.G.-A., T.A.M.P., M.A.Z., A.E.-M., J.B., V.C.-A., J.F., G.K., A.T., M.-J.S. and P.R.B. contributed to the data curation. M.B.-H, J.A., C.G.-A, M.R.-R, A.V.-C. and P.R.B designed the methodology, M.B.H., J.A., C.G.-A. and P.R.B. performed the analyses and M.B.-H, J.A. and P.R.B. performed the visualisation. T.A.M.P. and P.R.B. carried out the funding acquisition. J.A., S.S., and P.R.B. provided supervision and M.B.H, J.A., C.G.-A., M.R.-R and P.R.B. provided validation. M.B.H. and P.R.B. wrote the original draft of the manuscript. All authors contributed critically to the drafts and gave final approval for publication.

## 7. Data availability statement

The forest inventory data supporting this research are available from the original data producers, following the individual access policies, restrictions and licensing of each data owner. Open access to data is available for France (IGN, https://www.ign.fr/), Spain (Inventario Forestal Nacional, https://www.miteco.gob.es/) and the Netherlands (Dutch National Forest Inventory, https://www.wur.nl/). For other countries the data is available with an agreement from the data owners: the Polish National Forest Inventory and the Swedish National Forest Inventory (Swedish University of Agricultural Sciences). To facilitate the replication of our analyses, we have made a data and code repository available (https://doi.org/10.5281/zenodo.18836544). This repository includes all the necessary information for replication of the models, cross-validation and main figures. For confidentiality purposes, the plot coordinates are slightly perturbed (fuzzed) and the id of the original data was removed.

