## Supporting Information for "Contrasting trends in forest growth and mortality of major European tree species under increasing climatic stress"

**FOR ONLINE PUBLICATIONS ONLY**

**SUPPORTING INFORMATION**

### Appendix S1. Further details of the individual National Forest Inventories and tree growth and mortality variables.

**Table S1.1.** Main characteristics of the plot and sampling design of the National Forest Inventories (NFIs) as well as the number of species, plots and trees analysed in the study.

| **Country** | **Plot type** | **Mean plot area (ha)** | **First NFI measurement**  **[mean,**  **range]** | **Second NFI measurement**  **[mean,**  **range]** | **Mean census interval** | **No. species** | **No. target species** | **No. plots** | **No. trees** | **No. target trees** |
| --- | --- | --- | --- | --- | --- | --- | --- | --- | --- | --- |
| France | Concentric circles | 0.07 | 2012  (2010-2014) | 2017  (2015-2019) | 5 | 127 | 21 | 21,715 | 239,444 | 208,440 |
| Netherlands | Variable radius | 0.04 | 2004  (2001-2006) | 2013  (2013-2014) | 9.39 | 38 | 11 | 353 | 5,304 | 4,082 |
| Poland | Fixed radius | 0.03 | 2008  (2005-2010) | 2013  (2006-2015) | 4.98 | 65 | 14 | 10,922 | 200,196 | 181,418 |
| Spain | Concentric circles | 0.13 | 2002  (1997-2005) | 2016  (2005-2019) | 13.43 | 94 | 19 | 8,141 | 132,674 | 121,282 |
| Sweden | Concentric circles | 0.03 | 2011  (2008-2013) | 2015  (2013-2018) | 4.99 | 24 | 11 | 11,572 | 227,458 | 191,130 |
| **Total** |  |  |  |  |  |  |  | **52,703** | **805,076** | **706,352** |

**Table S1.2.** For each species, the country of occurrence, the total number of plots included in the study, and the number of plots with mortality occurrence (No. mortality, and the percentage of the total plot number). Mean species growth and mortality (m^2^ ha^-1^ yr^-1^) are also reported together with the 95% percentile range.

| **Species** | **Country** | **Total No. Plots**  **[No. mortality, %]** | **Growth (m^2^ ha^-1^ yr^-1^)** | **Mortality (m^2^ ha^-1^ yr^-1^)** |
| --- | --- | --- | --- | --- |
| *Abies alba* | France | 1840 [145, 5.49] | 0.563 [0.059, 1.485] | 0.56 [0.155, 1.29] |
|  | Poland | 799 [66, 2.5] | 0.593 [0.03, 1.531] | 0.403 [0.049, 1.278] |
|  | **Total** | **2639 [211, 7.99]** | **0.572 [0.047, 1.501]** | **0.041 [0, 0.325]** |
| *Alnus glutinosa* | France | 636 [91, 4.43] | 0.42 [0.022, 1.545] | 0.599 [0.18, 1.437] |
|  | Poland | 1078 [209, 10.17] | 0.449 [0.012, 1.342] | 0.313 [0.066, 0.815] |
|  | Sweden | 342 [37, 1.8] | 0.166 [0.006, 0.638] | 0.2 [0.055, 0.557] |
|  | **Total** | **2056 [337, 16.4]** | **0.393 [0.011, 1.328]** | **0.062 [0, 0.441]** |
| *Betula pendula* | France | 1839 [164, 3.09] | 0.224 [0.01, 0.817] | 0.444 [0.153, 0.988] |
|  | Poland | 3461 [346, 6.53] | 0.216 [0.007, 0.73] | 0.272 [0.046, 0.833] |
|  | **Total** | **5300 [510, 9.62]** | **0.219 [0.009, 0.753]** | **0.031 [0, 0.221]** |
| *Betula pubescens* | Sweden | 1471 [52, 3.54] | 0.036 [0.002, 0.114] | 0.193 [0.054, 0.472] |
|  | **Total** | **1471 [52, 3.54]** | **0.036 [0.002, 0.114]** | **0.007 [0, 0]** |
| *Carpinus betulus* | France | 3885 [164, 3.39] | 0.266 [0.026, 0.728] | 0.302 [0.144, 0.642] |
|  | Poland | 949 [48, 0.99] | 0.166 [0.011, 0.492] | 0.163 [0.04, 0.519] |
|  | **Total** | **4834 [212, 4.38]** | **0.246 [0.02, 0.698]** | **0.012 [0, 0]** |
| *Castanea sativa* | France | 2742 [650, 21.54] | 0.442 [0.026, 1.466] | 0.578 [0.153, 1.49] |
|  | Spain | 276 [95, 3.15] | 0.149 [0.008, 0.471] | 0.307 [0.047, 0.895] |
|  | **Total** | **3018 [745, 24.69]** | **0.416 [0.021, 1.386]** | **0.134 [0, 0.79]** |
| *Fagus sylvatica* | France | 4623 [198, 2.88] | 0.287 [0.033, 0.8] | 0.503 [0.162, 1.213] |
|  | Poland | 1619 [136, 1.98] | 0.372 [0.032, 1.021] | 0.36 [0.046, 1.283] |
|  | Spain | 639 [210, 3.05] | 0.144 [0.011, 0.374] | 0.208 [0.046, 0.568] |
|  | **Total** | **6881 [544, 7.91]** | **0.294 [0.028, 0.83]** | **0.028 [0, 0.164]** |
| *Fraxinus excelsior* | France | 2921 [232, 7.25] | 0.261 [0.023, 0.826] | 0.581 [0.162, 1.573] |
|  | Poland | 280 [70, 2.19] | 0.14 [0.005, 0.465] | 0.54 [0.055, 1.746] |
|  | **Total** | **3201 [302, 9.44]** | **0.25 [0.02, 0.801]** | **0.054 [0, 0.38]** |
| *Picea abies* | France | 1782 [188, 1.54] | 0.561 [0.041, 1.704] | 0.621 [0.162, 1.544] |
|  | Poland | 2331 [446, 3.65] | 0.38 [0.01, 1.324] | 0.334 [0.048, 1.383] |
|  | Sweden | 8112 [1048, 8.57] | 0.349 [0.012, 1.151] | 0.29 [0.054, 0.918] |
|  | **Total** | **12225 [1682, 13.76]** | **0.386 [0.013, 1.308]** | **0.047 [0, 0.277]** |
| *Pinus halepensis* | Spain | 1267 [355, 28.02] | 0.101 [0.009, 0.282] | 0.167 [0.037, 0.526] |
|  | **Total** | **1267 [355, 28.02]** | **0.101 [0.009, 0.282]** | **0.047 [0, 0.257]** |
| *Pinus nigra* | Spain | 723 [129, 17.84] | 0.121 [0.007, 0.381] | 0.134 [0.034, 0.374] |
|  | **Total** | **723 [129, 17.84]** | **0.121 [0.007, 0.381]** | **0.024 [0, 0.132]** |
| *Pinus pinaster* | France | 1216 [126, 5.52] | 0.706 [0.052, 2.24] | 0.549 [0.153, 1.057] |
|  | Spain | 1065 [390, 17.1] | 0.156 [0.009, 0.519] | 0.223 [0.039, 0.664] |
|  | **Total** | **2281 [516, 22.62]** | **0.45 [0.017, 1.801]** | **0.069 [0, 0.47]** |
| *Pinus pinea* | Spain | 439 [89, 20.27] | 0.103 [0.009, 0.288] | 0.162 [0.031, 0.54] |
|  | **Total** | **439 [89, 20.27]** | **0.103 [0.009, 0.288]** | **0.033 [0, 0.191]** |
| *Pinus sylvestris* | France | 2686 [307, 1.55] | 0.309 [0.018, 0.969] | 0.521 [0.162, 1.209] |
|  | Netherlands | 212 [74, 0.37] | 0.477 [0.047, 1.623] | 0.271 [0.042, 0.719] |
|  | Poland | 7343 [1060, 5.34] | 0.706 [0.085, 1.715] | 0.309 [0.082, 0.9] |
|  | Spain | 1342 [546, 2.75] | 0.165 [0.009, 0.503] | 0.197 [0.039, 0.616] |
|  | Sweden | 8250 [944, 4.76] | 0.282 [0.022, 0.875] | 0.23 [0.053, 0.75] |
|  | **Total** | **19833 [2931, 14.77]** | **0.437 [0.024, 1.313]** | **0.042 [0, 0.282]** |
| *Populus tremula* | France | 926 [106, 5.35] | 0.34 [0.028, 1.155] | 0.45 [0.155, 0.805] |
|  | Poland | 490 [61, 3.08] | 0.274 [0.011, 0.845] | 0.293 [0.058, 0.841] |
|  | Sweden | 564 [40, 2.02] | 0.162 [0.006, 0.609] | 0.22 [0.053, 0.861] |
|  | **Total** | **1980 [207, 10.45]** | **0.273 [0.012, 0.932]** | **0.038 [0, 0.324]** |
| *Quercus ilex* | France | 938 [32, 0.96] | 0.308 [0.021, 0.829] | 0.276 [0.144, 0.536] |
|  | Spain | 2381 [247, 7.44] | 0.058 [0.005, 0.171] | 0.105 [0.032, 0.294] |
|  | **Total** | **3319 [279, 8.4]** | **0.129 [0.006, 0.527]** | **0.01 [0, 0.067]** |
| *Quercus petraea* | France | 4722 [312, 6.26] | 0.272 [0.028, 0.775] | 0.397 [0.153, 0.83] |
|  | Spain | 264 [62, 1.24] | 0.089 [0.005, 0.308] | 0.132 [0.037, 0.366] |
|  | **Total** | **4986 [374, 7.5]** | **0.262 [0.022, 0.75]** | **0.026 [0, 0.203]** |
| *Quercus pubescens* | France | 2866 [222, 6.81] | 0.266 [0.022, 0.762] | 0.334 [0.144, 0.799] |
|  | Spain | 394 [41, 1.26] | 0.052 [0.005, 0.18] | 0.092 [0.032, 0.206] |
|  | **Total** | **3260 [263, 8.07]** | **0.24 [0.014, 0.734]** | **0.024 [0, 0.193]** |
| *Quercus pyrenaica* | Spain | 1154 [326, 28.25] | 0.093 [0.007, 0.262] | 0.128 [0.033, 0.337] |
|  | **Total** | **1154 [326, 28.25]** | **0.093 [0.007, 0.262]** | **0.036 [0, 0.19]** |
| *Quercus robur* | France | 6530 [504, 7] | 0.251 [0.023, 0.74] | 0.482 [0.153, 1.149] |
|  | Netherlands | 225 [84, 1.17] | 0.308 [0.02, 1.07] | 0.226 [0.04, 0.613] |
|  | Spain | 448 [85, 1.18] | 0.103 [0.006, 0.317] | 0.208 [0.046, 0.52] |
|  | **Total** | **7203 [673, 9.35]** | **0.243 [0.02, 0.725]** | **0.039 [0, 0.325]** |
| *Salix caprea* | France | 622 [138, 22.19] | 0.348 [0.013, 1.251] | 0.458 [0.153, 1.197] |
|  | **Total** | **622 [138, 22.19]** | **0.348 [0.013, 1.251]** | **0.102 [0, 0.611]** |

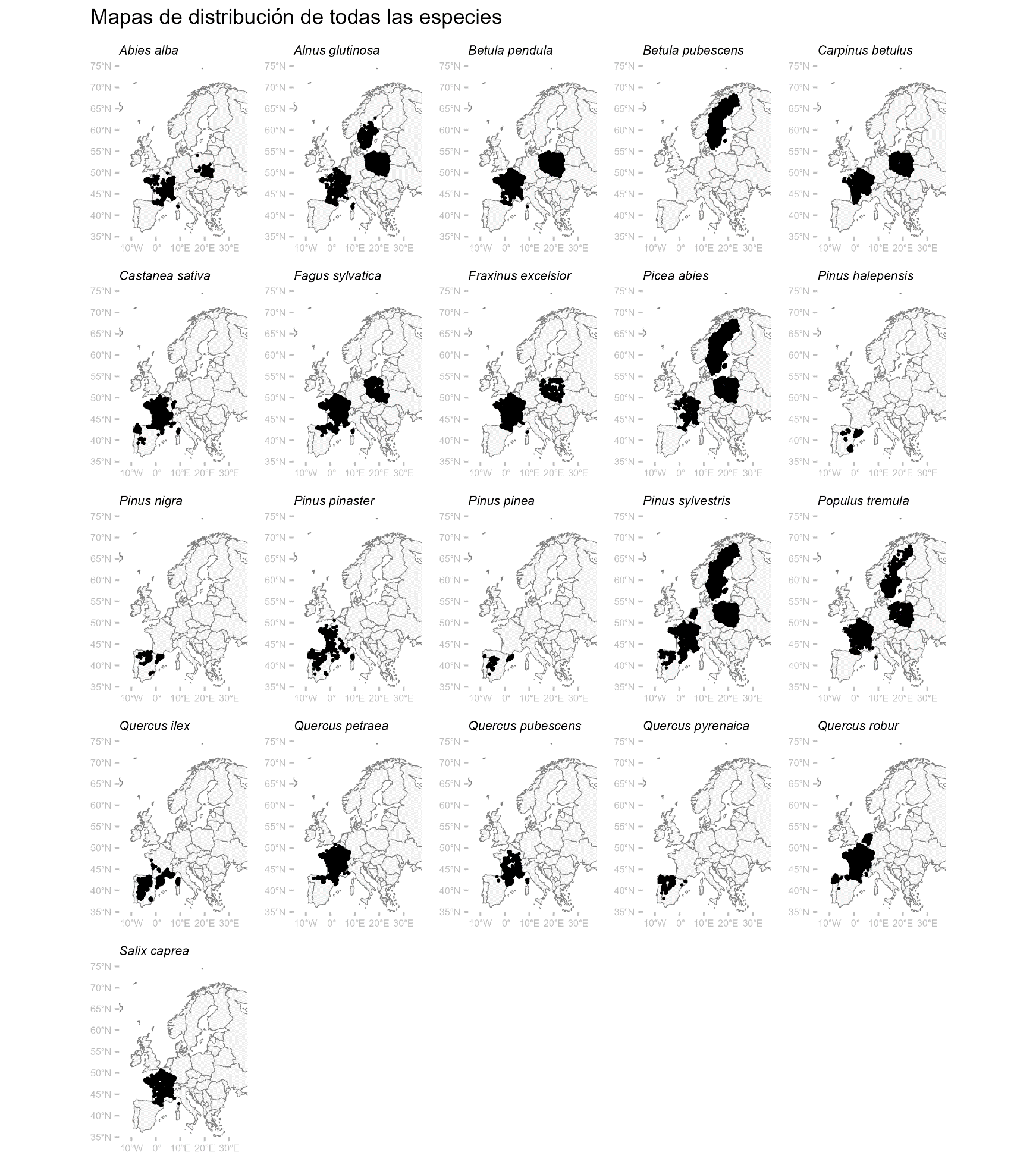

**Figure S1.1.** Species-specific NFI plots used in the analyses and their spatial distribution.

### **Appendix S2.** Further details on variable selection, explanatory variable characteristics and data coverage.

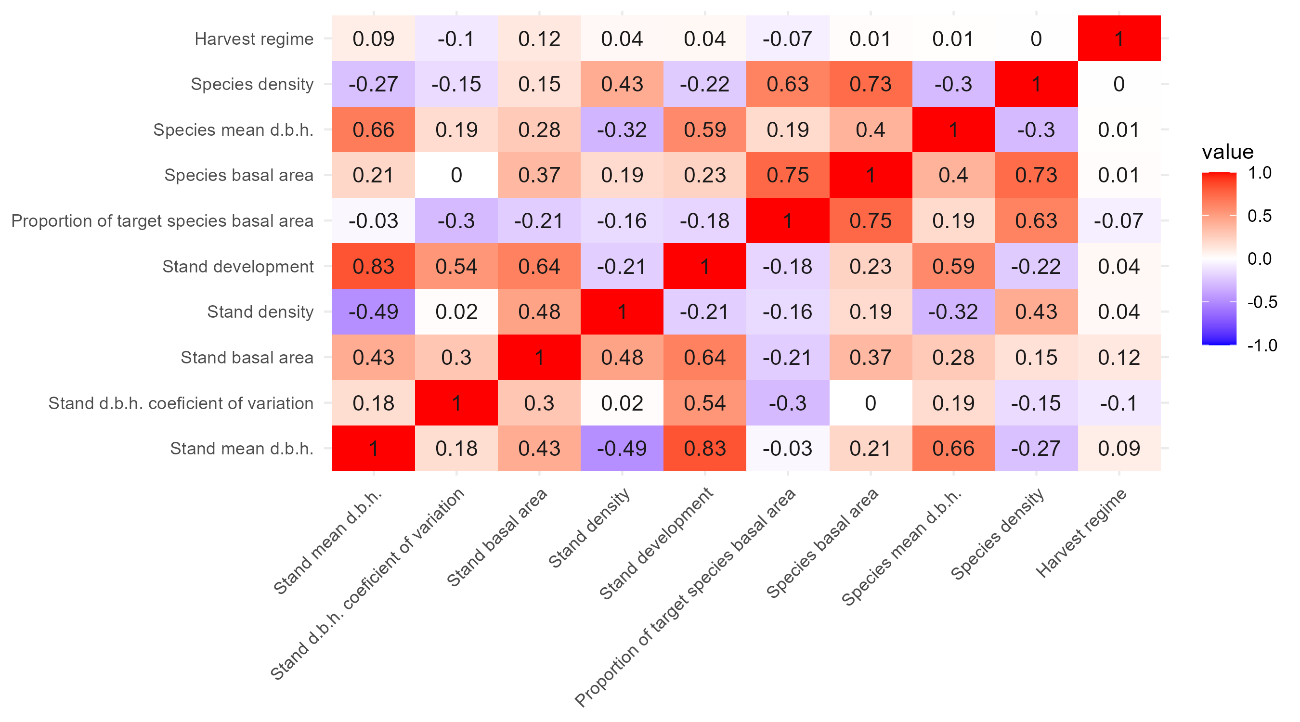

**Figure S2.1.** Heatmap of correlations between all initially calculated structural variables showing Spearman coefficient (rho) considering all plots and species.

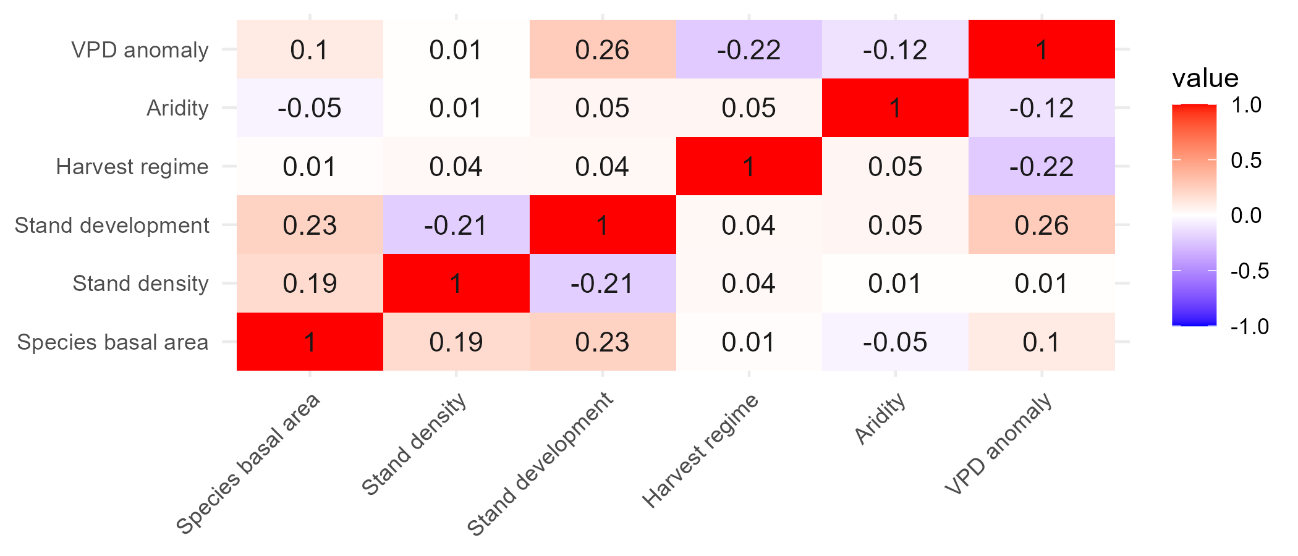

**Figure S2.2.** Heatmap of correlations between the explanatory variables included in the models showing Spearman coefficient (rho) considering all plots and species.

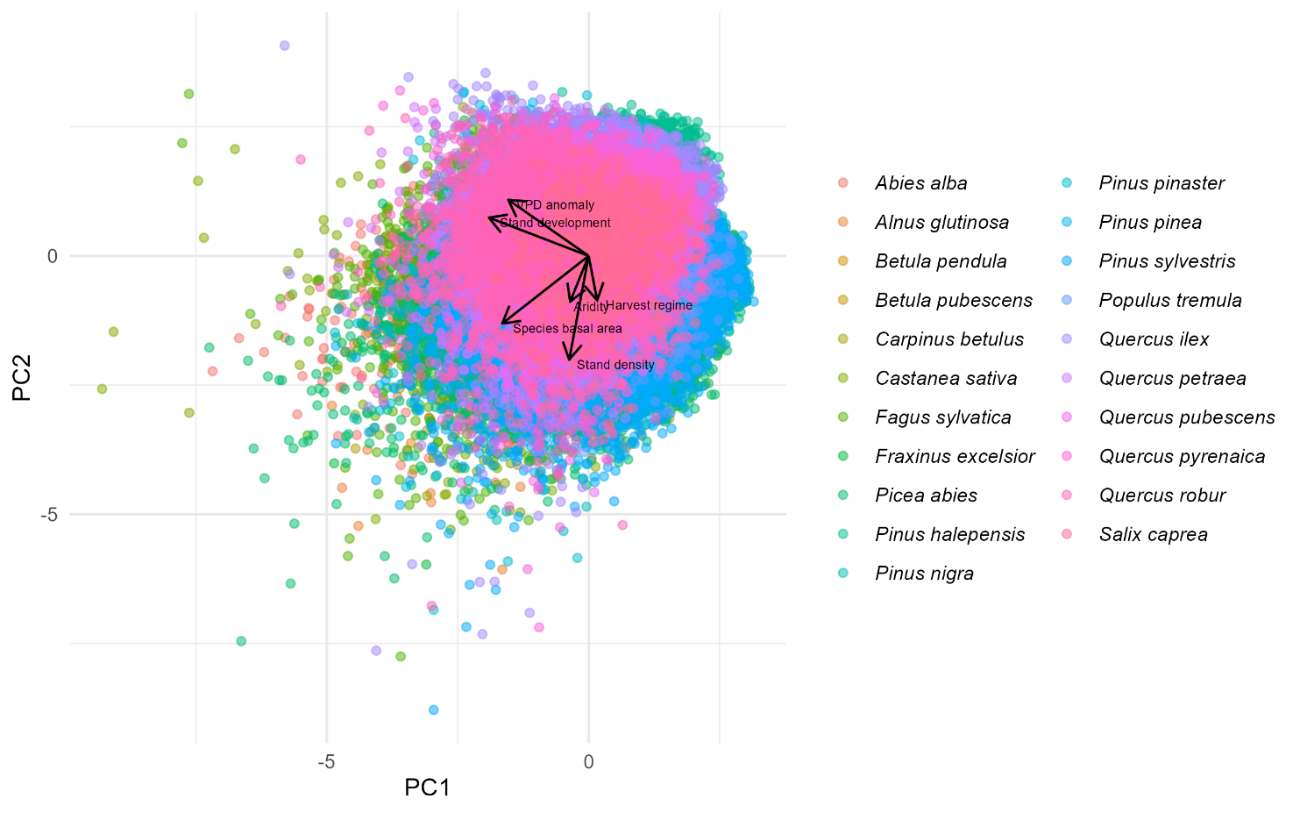

**Figure S2.3.** Principal components analyses (PCA) for the explanatory variables included in the models considering all plots and species.

**Table S2.1.** Correlation between explanatory variables included in the models showing Spearman coefficient (rho) for all target species. Bold values indicate correlations equal to or greater than 0.5. Highest correlation value was 0.7.

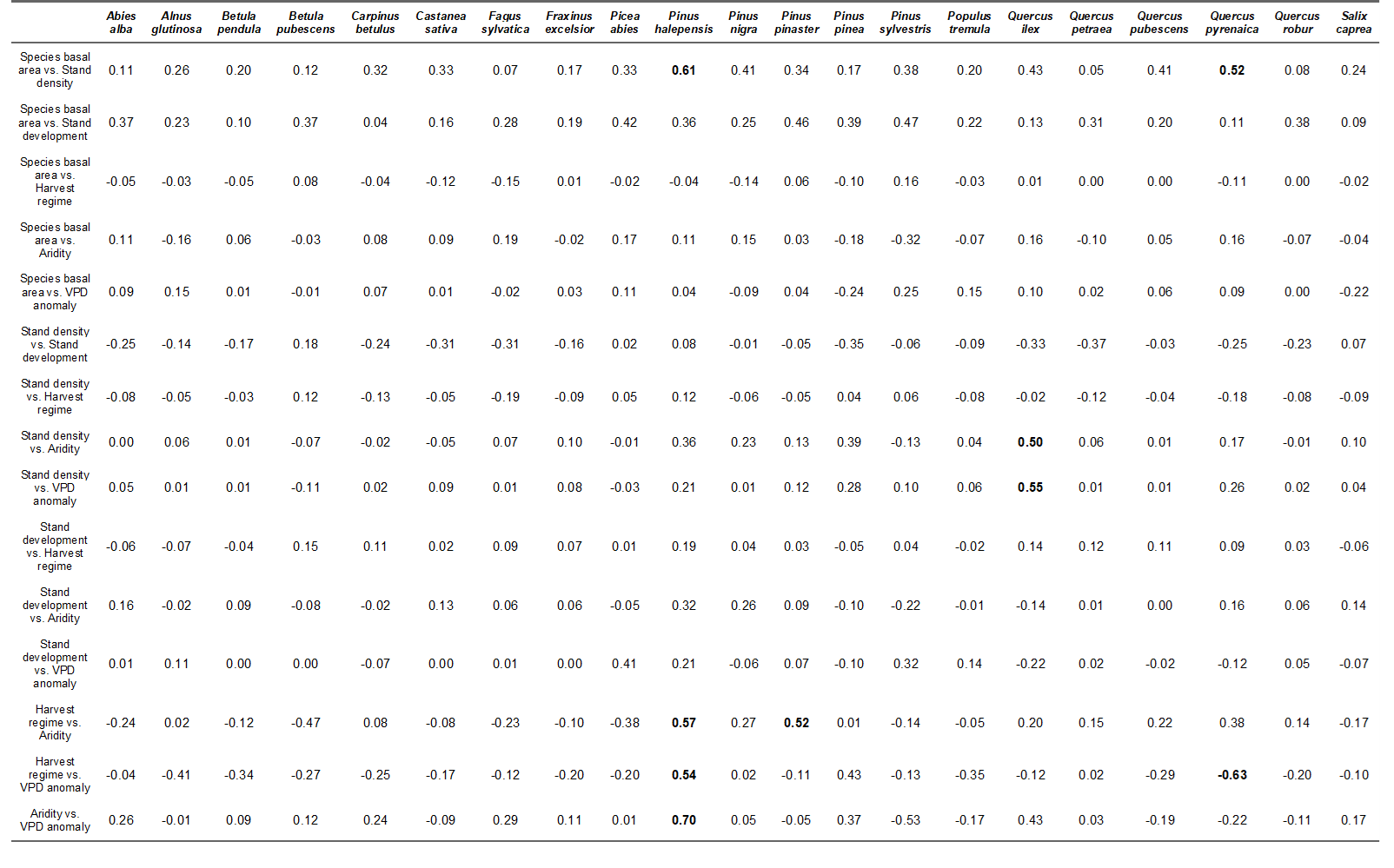

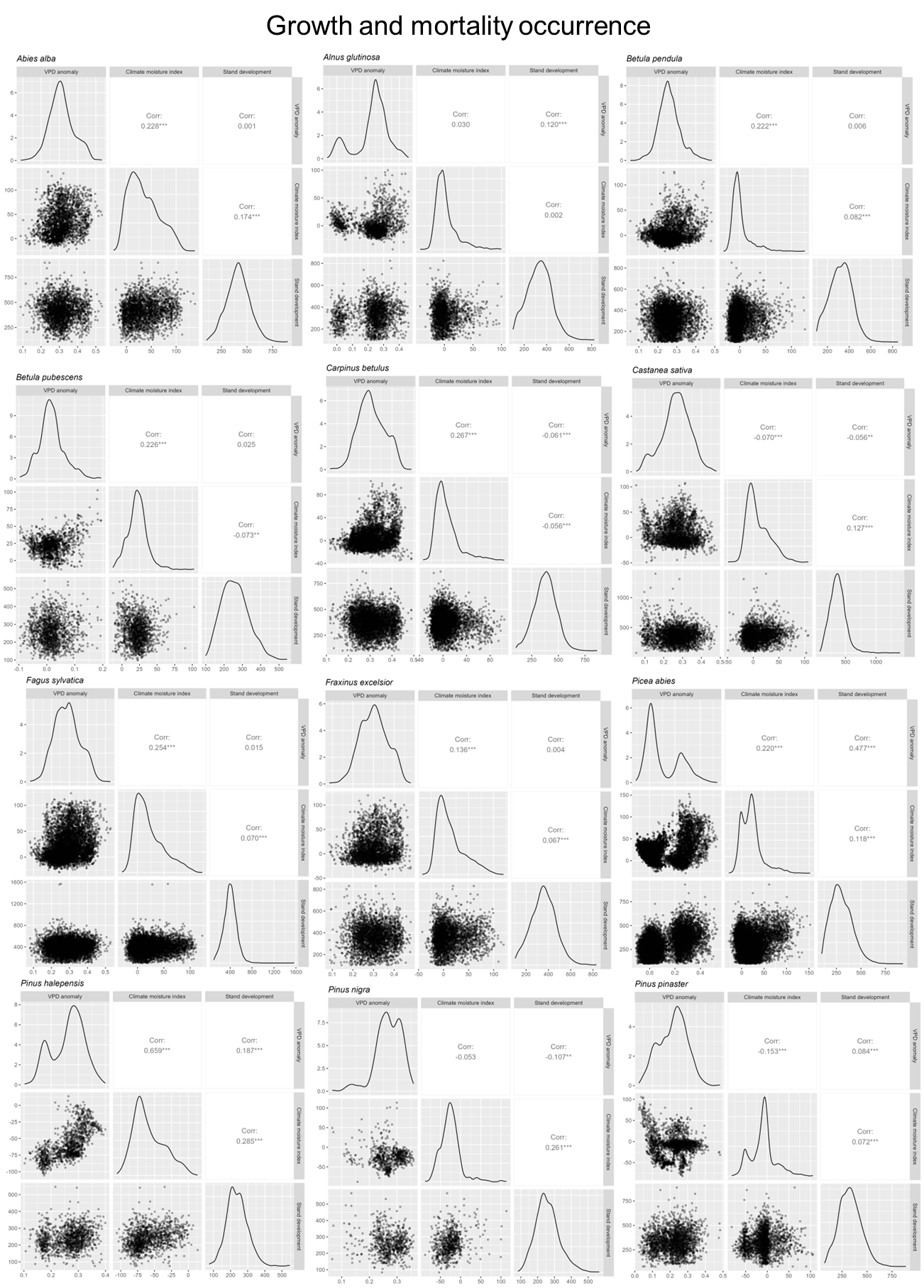

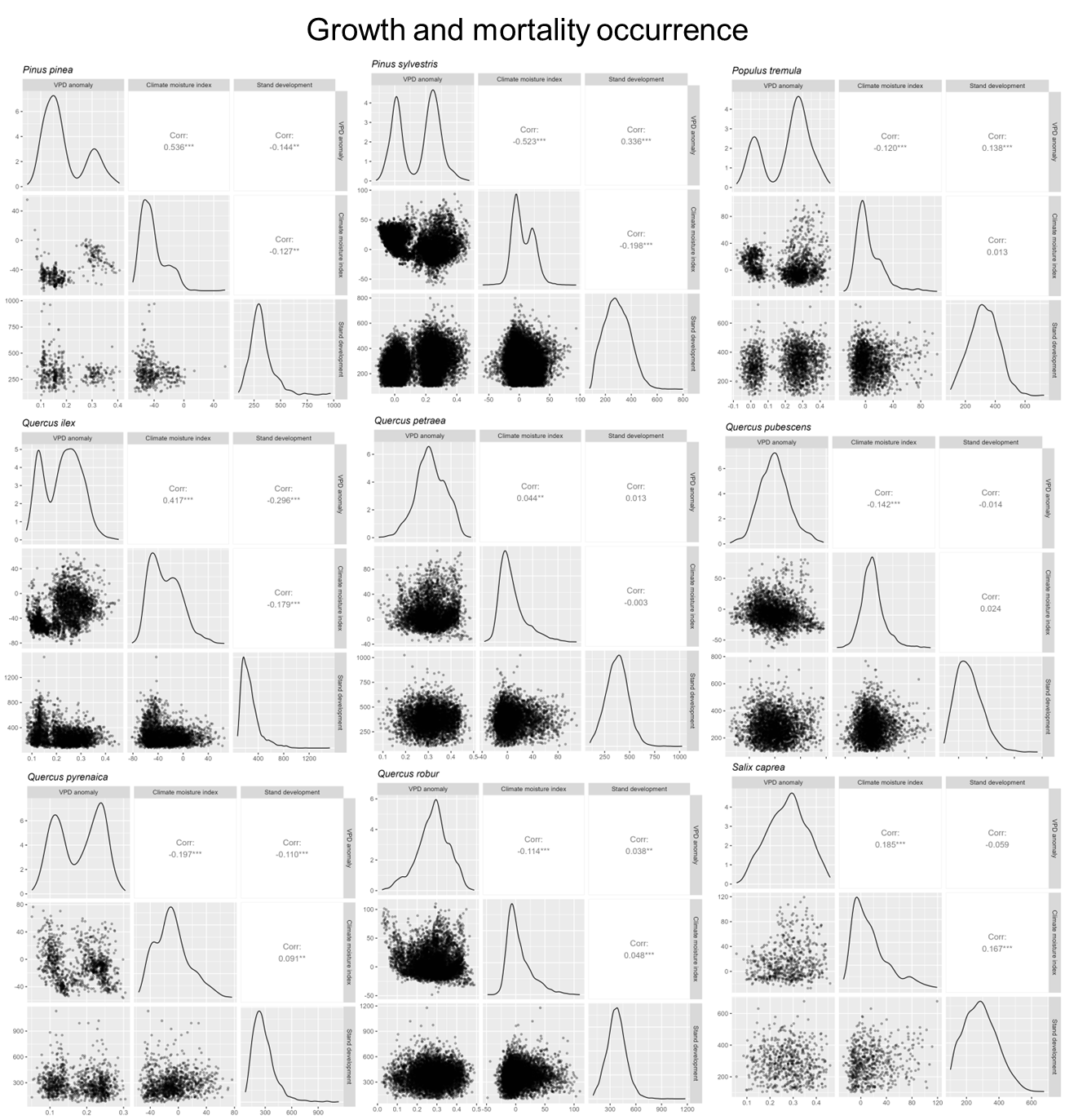

**Figure S2.4.** Data coverage across the full three-way interaction space for growth and mortality occurrence responses.

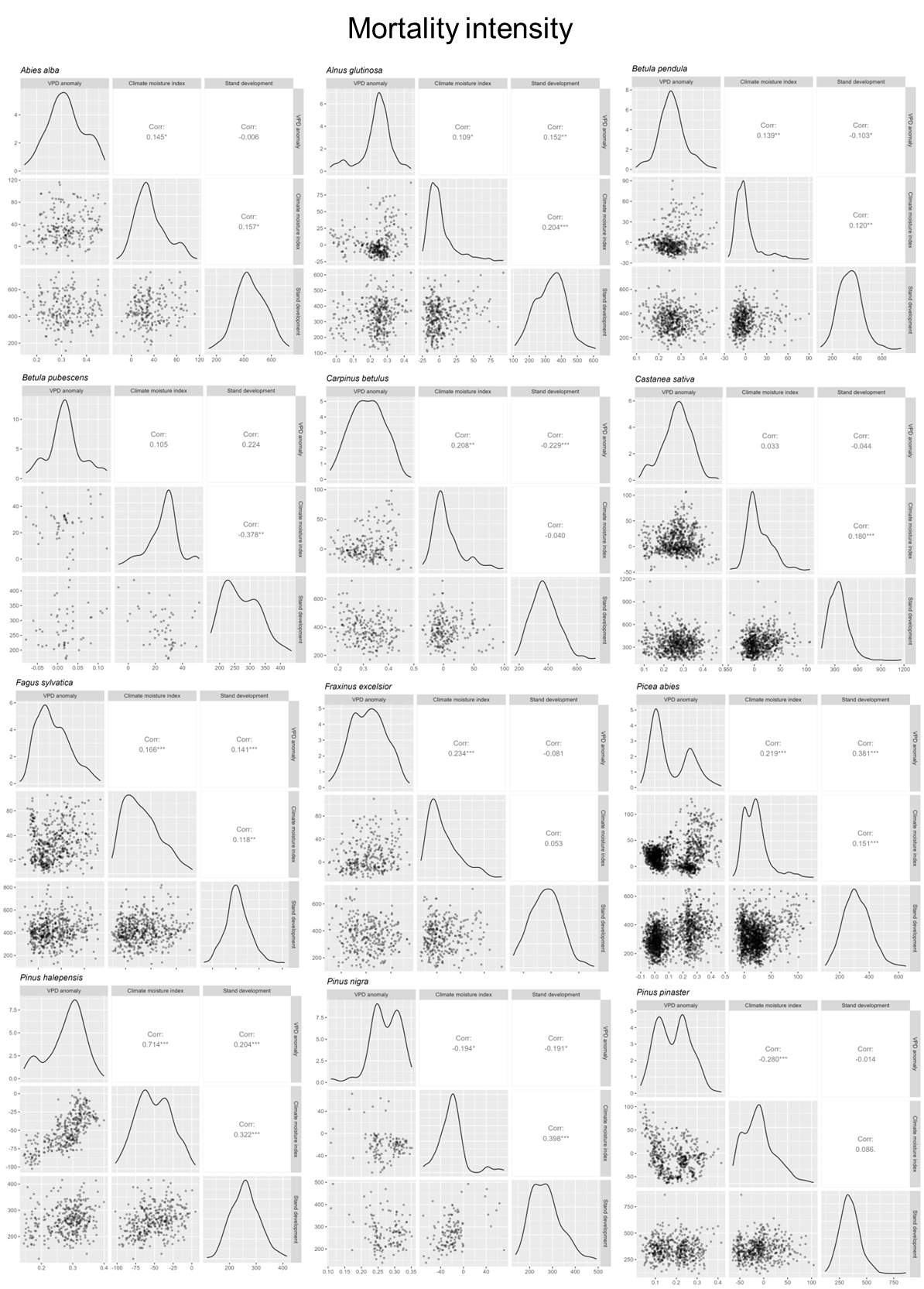

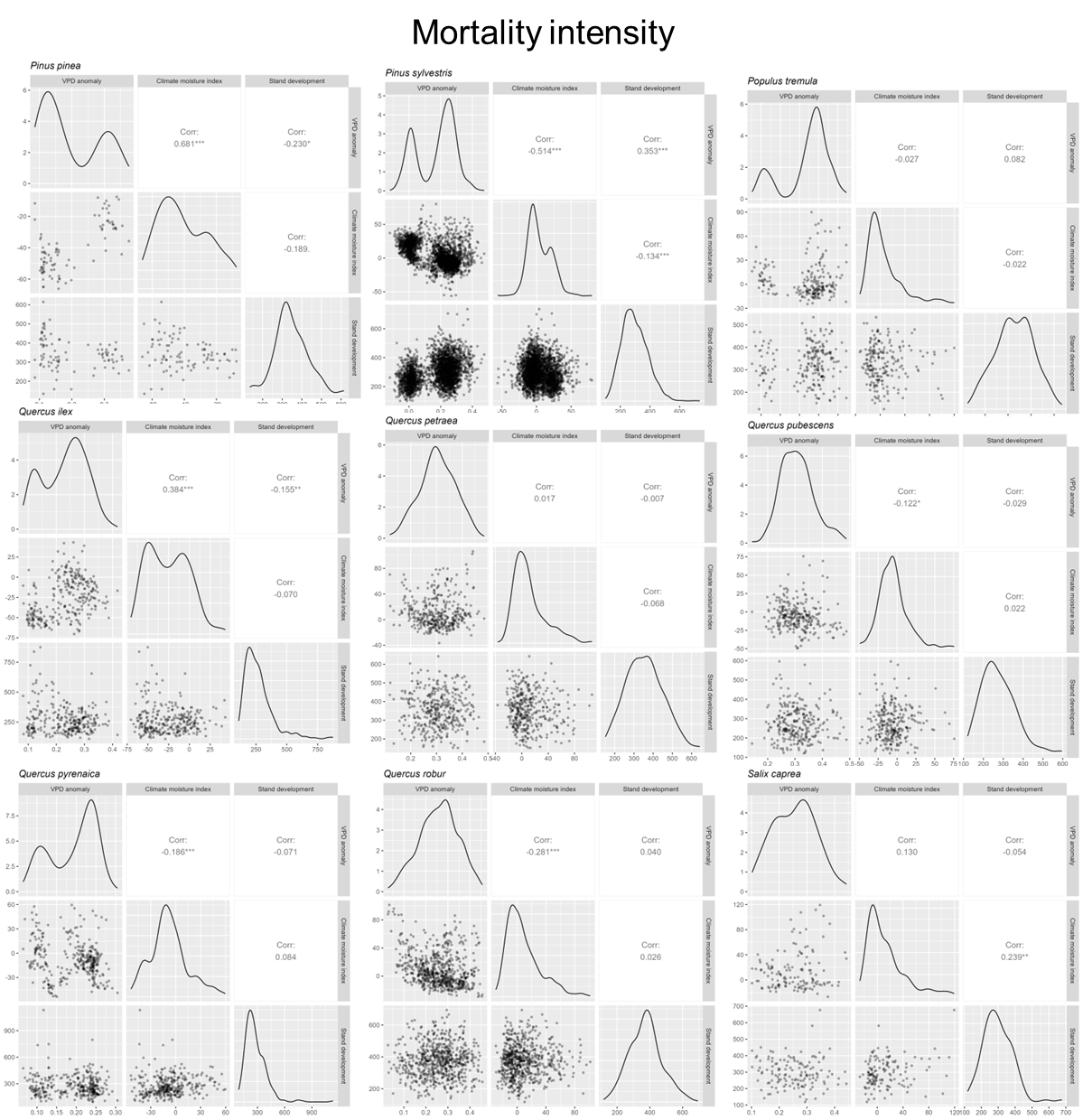

**Figure S2.5.** Data coverage across the full three-way interaction space for mortality intensity responses.

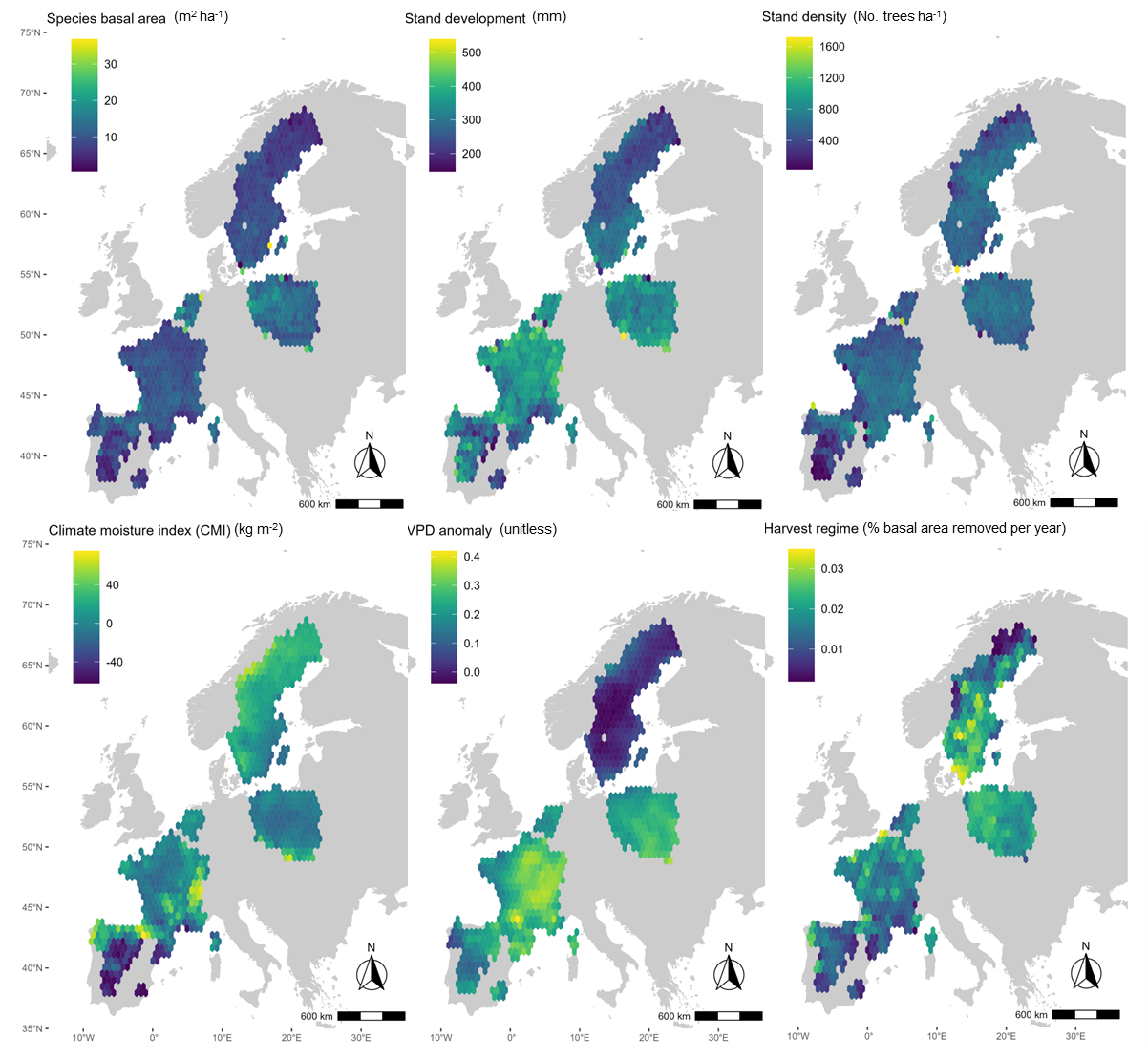

**Figure S2.5.** Spatial patterns of mean explanatory variables across the study area, aggregated to a coarser spatial resolution (c. 0.6º x 0.6º latitude-longitude grid cells) than that used in the models to enhance visualisation.

### Appendix S3. Further details on model validation.

**Table S3.1.** Explained deviance (Expl. Dev.) for species-level growth and mortality models (occurrence and intensity), including the significance for the triple interaction between stand development, aridity and VPD anomaly. Significant values are highlighted (P-value < 0.05).

| **Species** | **Growth** | | **Mortality ocurrence** | | **Mortality intensity** | |
| --- | --- | --- | --- | --- | --- | --- |
|  | **Expl. Dev.** | **Interaction**  **P-value** | **Expl. Dev.** | **Interaction**  **P-value** | **Expl. Dev.** | **Interaction**  **P-value** |
| *Abies alba* | 0.561 | **<0.001** | 0.135 | 0.9 | 0.249 | 0.51 |
| *Alnus glutinosa* | 0.578 | **<0.001** | 0.165 | **0.003** | 0.358 | 0.425 |
| *Betula pendula* | 0.438 | **<0.001** | 0.146 | **<0.001** | 0.273 | 0.799 |
| *Betula pubescens* | 0.219 | **0.001** | 0.318 | 0.325 | 0.595 | 0.183 |
| *Carpinus betulus* | 0.307 | **<0.001** | 0.168 | **<0.001** | 0.305 | **0.037** |
| *Castanea sativa* | 0.445 | **<0.001** | 0.237 | **<0.001** | 0.250 | **0.032** |
| *Fagus sylvatica* | 0.448 | **<0.001** | 0.266 | 0.211 | 0.309 | 0.269 |
| *Fraxinus excelsior* | 0.458 | **<0.001** | 0.197 | **<0.001** | 0.352 | **0.007** |
| *Picea abies* | 0.468 | **<0.001** | 0.152 | **<0.001** | 0.359 | **<0.001** |
| *Pinus halepensis* | 0.358 | **<0.001** | 0.278 | **<0.001** | 0.305 | **0.018** |
| *Pinus nigra* | 0.531 | **<0.001** | 0.173 | 0.78 | 0.331 | 0.848 |
| *Pinus pinaster* | 0.682 | **<0.001** | 0.304 | **0.009** | 0.456 | **0.001** |
| *Pinus pinea* | 0.477 | **<0.001** | 0.229 | **0.005** | 0.376 | 0.412 |
| *Pinus sylvestris* | 0.588 | **<0.001** | 0.154 | **<0.001** | 0.265 | **<0.001** |
| *Populus tremula* | 0.563 | **<0.001** | 0.171 | 0.065 | 0.279 | 0.331 |
| *Quercus ilex* | 0.623 | **<0.001** | 0.151 | **0.025** | 0.302 | 0.578 |
| *Quercus petraea* | 0.557 | **<0.001** | 0.190 | **0.003** | 0.365 | **0.014** |
| *Quercus pubescens* | 0.480 | **<0.001** | 0.142 | 0.168 | 0.451 | 0.262 |
| *Quercus pyrenaica* | 0.443 | **0.001** | 0.235 | 0.202 | 0.187 | 0.148 |
| *Quercus robur* | 0.511 | **<0.001** | 0.209 | **<0.001** | 0.298 | **0.031** |
| *Salix caprea* | 0.413 | **<0.001** | 0.252 | 0.076 | 0.460 | 0.59 |

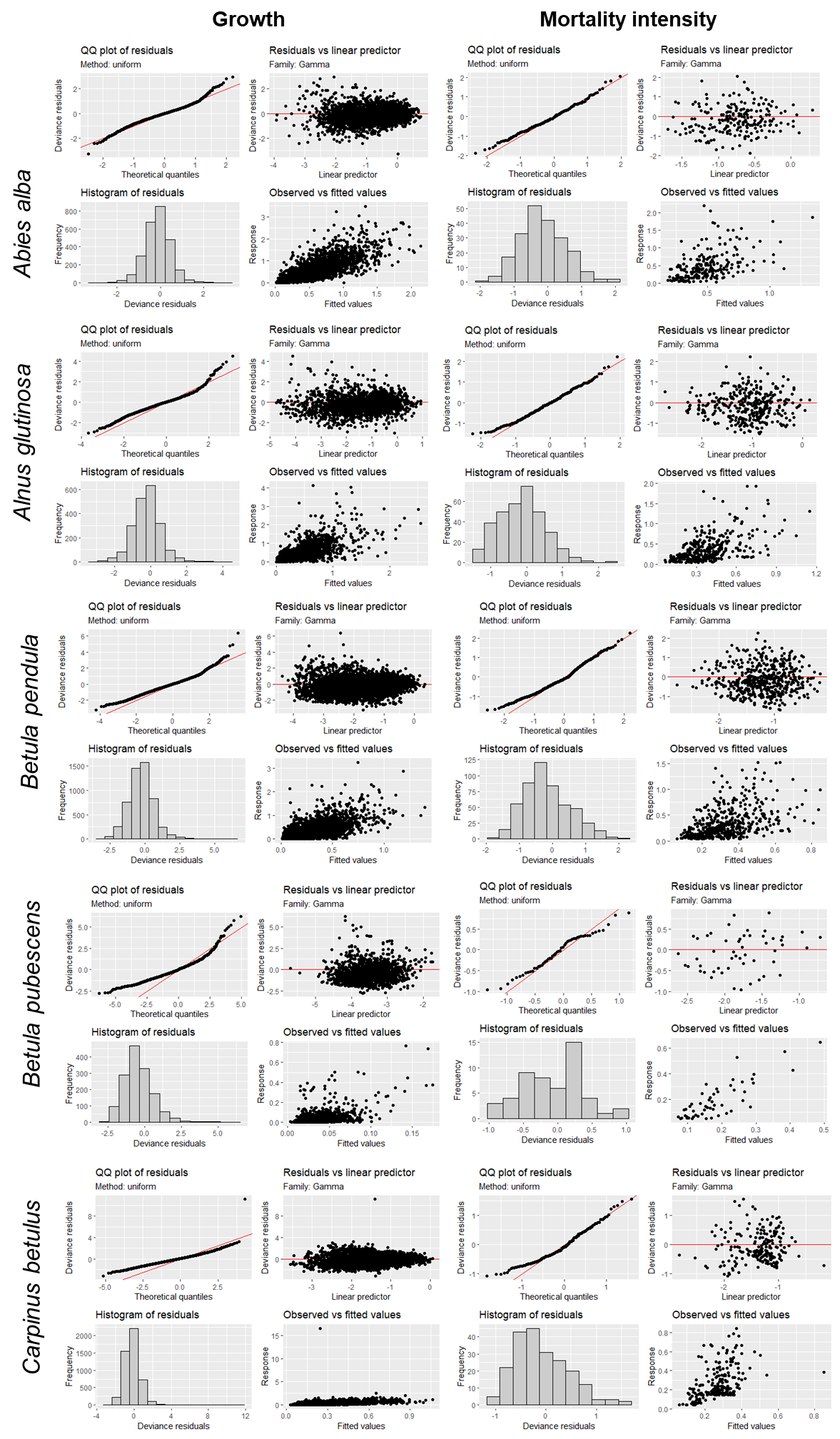

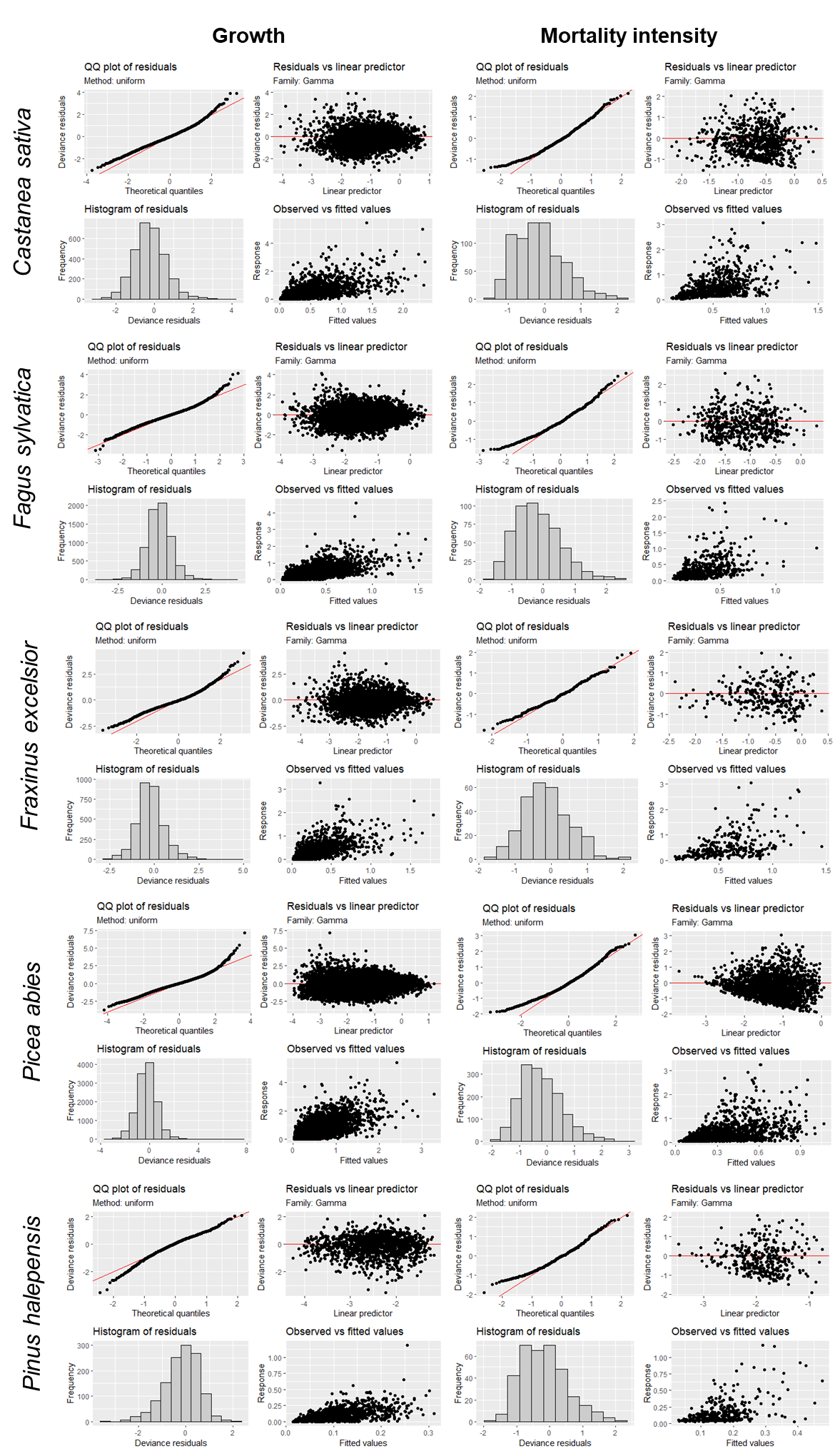

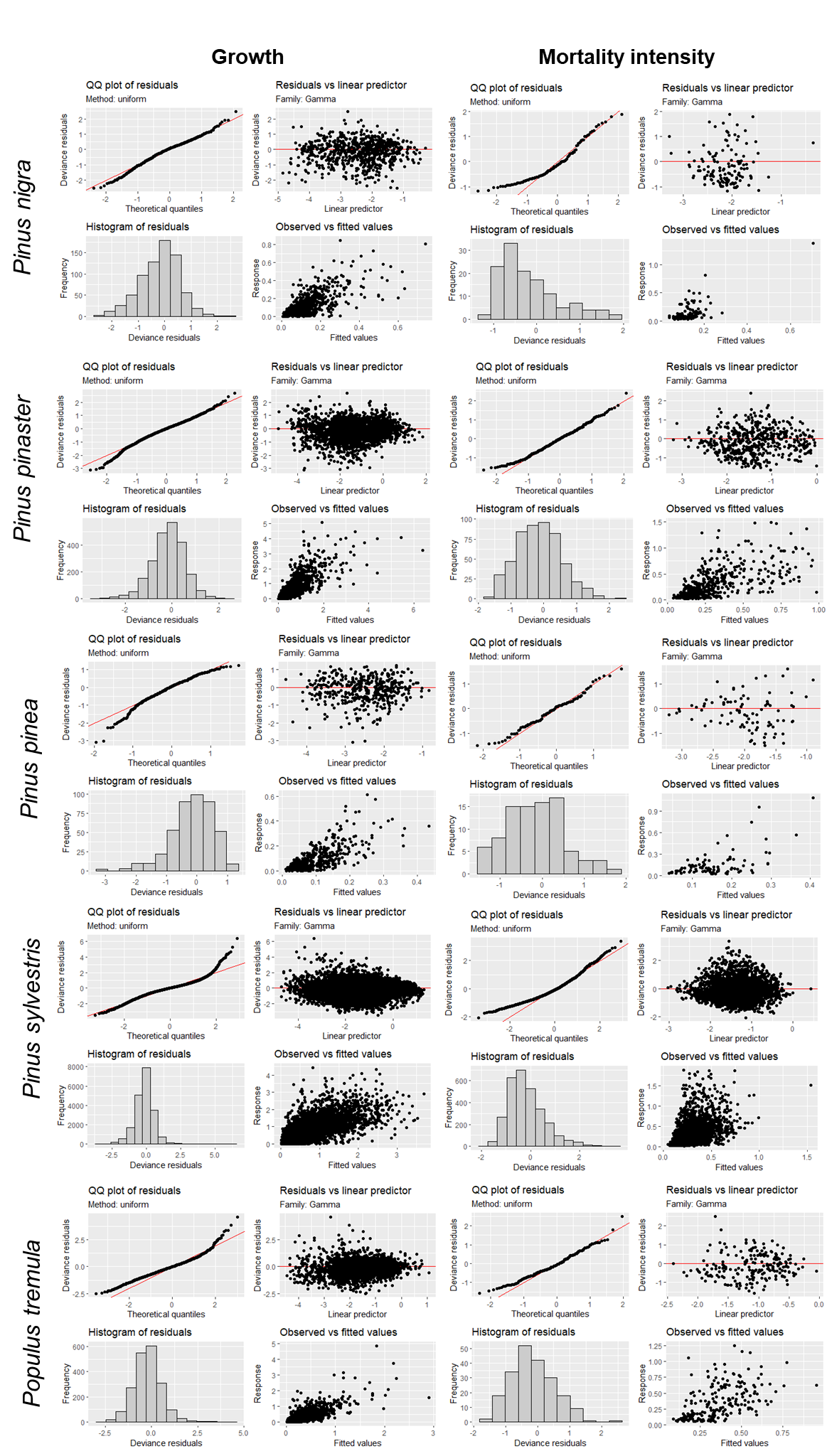

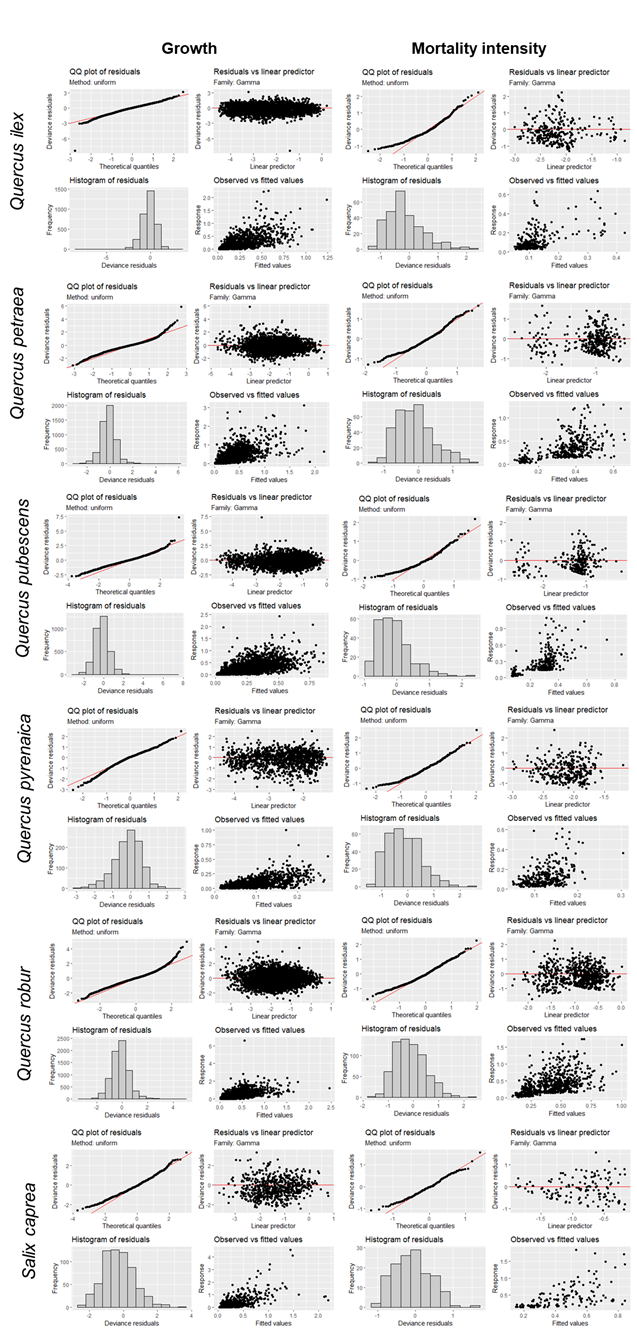

**Figure S3.1.** Model diagnostics showing species-specific models residuals for growth and mortality intensity models.

**Table S3.2.** Species growth and mortality intensity model goodness of fit. Mean predictive performance ± standard deviation is represented based on: R-squared, Root Mean Square Error (RMSE) and Mean Absolute Error (MAE) and their normalised values, NRMSE and NMAE, respectively. Mean values per species correspond to the mean of the 500 iterations.

| **Growth** | | | | |
| --- | --- | --- | --- | --- |
| **Species** | **RMSE** | **NMRSE** | **MAE** | **NMAE** |
| *Abies alba* | 0.314 ± 0.017 | 0.508 | 0.217 ± 0.009 | 0.351 |
| *Alnus glutinosa* | 0.334 ± 0.034 | 0.700 | 0.191 ± 0.011 | 0.400 |
| *Betula pendula* | 0.197 ± 0.013 | 0.833 | 0.12 ± 0.004 | 0.506 |
| *Betula pubescens* | 0.056 ± 0.009 | 1.705 | 0.029 ± 0.002 | 0.871 |
| *Carpinus betulus* | 0.272 ± 0.156 | 1.089 | 0.134 ± 0.007 | 0.537 |
| *Castanea sativa* | 0.382 ± 0.029 | 0.856 | 0.233 ± 0.01 | 0.522 |
| *Fagus sylvatica* | 0.222 ± 0.051 | 0.742 | 0.138 ± 0.005 | 0.460 |
| *Fraxinus excelsior* | 0.217 ± 0.019 | 0.850 | 0.13 ± 0.006 | 0.509 |
| *Larix decidua* | 0.311 ± 0.011 | 0.729 | 0.196 ± 0.004 | 0.460 |
| *Picea abies* | 0.079 ± 0.01 | 0.745 | 0.054 ± 0.003 | 0.507 |
| *Pinus halepensis* | 0.092 ± 0.009 | 0.660 | 0.061 ± 0.005 | 0.438 |
| *Pinus nigra* | 0.364 ± 0.037 | 0.771 | 0.197 ± 0.013 | 0.418 |
| *Pinus pinaster* | 0.07 ± 0.009 | 0.680 | 0.049 ± 0.005 | 0.476 |
| *Pinus pinea* | 0.282 ± 0.008 | 0.573 | 0.173 ± 0.003 | 0.352 |
| *Pinus sylvestris* | 0.233 ± 0.029 | 0.822 | 0.133 ± 0.008 | 0.470 |
| *Populus tremula* | 0.14 ± 0.014 | 1.249 | 0.072 ± 0.004 | 0.641 |
| *Prunus avium* | 0.192 ± 0.014 | 0.718 | 0.112 ± 0.004 | 0.417 |
| *Quercus ilex* | 0.19 ± 0.012 | 0.684 | 0.122 ± 0.005 | 0.440 |
| *Quercus petraea* | 0.072 ± 0.01 | 0.683 | 0.048 ± 0.003 | 0.452 |
| *Quercus pubescens* | 0.196 ± 0.026 | 0.790 | 0.112 ± 0.004 | 0.454 |
| *Quercus pyrenaica* | 0.397 ± 0.073 | 1.070 | 0.232 ± 0.025 | 0.624 |
| *Quercus robur* | 0.314 ± 0.017 | 0.508 | 0.217 ± 0.009 | 0.351 |
| *Salix caprea* | 0.334 ± 0.034 | 0.700 | 0.191 ± 0.011 | 0.400 |

| **Mortality intensity** | | | | |
| --- | --- | --- | --- | --- |
| **Species** | **RMSE** | **NMRSE** | **MAE** | **NMAE** |
| *Abies alba* | 0.399+-0.064 | 0.937 | 0.295+-0.036 | 0.692 |
| *Alnus glutinosa* | 0.296+-0.041 | 0.818 | 0.207+-0.022 | 0.574 |
| *Betula pendula* | 0.265+-0.026 | 0.862 | 0.191+-0.015 | 0.619 |
| *Betula pubescens* | 0.367+-1.026 | 2.383 | 0.216+-0.445 | 1.401 |
| *Carpinus betulus* | 0.169+-0.025 | 0.836 | 0.13+-0.014 | 0.643 |
| *Castanea sativa* | 0.428+-0.039 | 0.910 | 0.301+-0.02 | 0.641 |
| *Fagus sylvatica* | 0.325+-0.044 | 1.012 | 0.216+-0.019 | 0.673 |
| *Fraxinus excelsior* | 0.487+-0.074 | 1.046 | 0.338+-0.041 | 0.726 |
| *Larix decidua* | 0.362+-0.033 | 1.151 | 0.224+-0.013 | 0.714 |
| *Picea abies* | 0.161+-0.026 | 1.254 | 0.105+-0.013 | 0.822 |
| *Pinus halepensis* | 0.265+-0.34 | 2.736 | 0.123+-0.075 | 1.268 |
| *Pinus nigra* | 0.236+-0.025 | 0.685 | 0.163+-0.015 | 0.474 |
| *Pinus pinaster* | 0.171+-0.058 | 1.445 | 0.116+-0.03 | 0.980 |
| *Pinus pinea* | 0.25+-0.014 | 0.948 | 0.167+-0.006 | 0.635 |
| *Pinus sylvestris* | 0.248+-0.033 | 0.641 | 0.191+-0.021 | 0.493 |
| *Populus tremula* | 0.102+-0.016 | 0.913 | 0.07+-0.008 | 0.630 |
| *Prunus avium* | 0.213+-0.024 | 0.770 | 0.155+-0.015 | 0.560 |
| *Quercus ilex* | 0.19+-0.032 | 0.874 | 0.133+-0.017 | 0.610 |
| *Quercus petraea* | 0.101+-0.015 | 0.970 | 0.071+-0.007 | 0.688 |
| *Quercus pubescens* | 0.27+-0.021 | 0.762 | 0.203+-0.013 | 0.573 |
| *Quercus pyrenaica* | 0.454+-0.827 | 1.304 | 0.27+-0.157 | 0.774 |
| *Quercus robur* | 0.399+-0.064 | 0.937 | 0.295+-0.036 | 0.692 |
| *Salix caprea* | 0.296+-0.041 | 0.818 | 0.207+-0.022 | 0.574 |

**Table S3.3.** Mean predictive performance of mortality occurrence models based on AUC (i.e. area under the Receiver Operating Characteristic – ROC curve) and Se* (i.e. probability of correctly classifying any given case, calculated as the average of sensitivity and specificity). Mean values per species obtained from the 500 iterations ± standard deviation are represented.

| **Mortality occurrence** | | | | |
| --- | --- | --- | --- | --- |
| **Species** | **AUC** | **Se*** | **Sensitivity** | **Specificity** |
| *Abies alba* | 0.758 ± 0.031 | 0.696 ± 0.027 | 0.974 ± 0.009 | 0.17 ± 0.033 |
| *Alnus glutinosa* | 0.766 ± 0.027 | 0.698 ± 0.027 | 0.936 ± 0.018 | 0.325 ± 0.049 |
| *Betula pendula* | 0.778 ± 0.02 | 0.707 ± 0.019 | 0.967 ± 0.008 | 0.201 ± 0.024 |
| *Betula pubescens* | 0.859 ± 0.05 | 0.77 ± 0.068 | 0.993 ± 0.006 | 0.151 ± 0.07 |
| *Carpinus betulus* | 0.796 ± 0.031 | 0.718 ± 0.034 | 0.987 ± 0.005 | 0.112 ± 0.028 |
| *Castanea sativa* | 0.814 ± 0.018 | 0.735 ± 0.019 | 0.905 ± 0.019 | 0.49 ± 0.049 |
| *Fagus sylvatica* | 0.845 ± 0.017 | 0.76 ± 0.021 | 0.974 ± 0.006 | 0.249 ± 0.046 |
| *Fraxinus excelsior* | 0.806 ± 0.025 | 0.729 ± 0.027 | 0.969 ± 0.01 | 0.232 ± 0.039 |
| *Larix decidua* | 0.771 ± 0.01 | 0.697 ± 0.011 | 0.943 ± 0.008 | 0.265 ± 0.022 |
| *Picea abies* | 0.827 ± 0.026 | 0.747 ± 0.028 | 0.895 ± 0.029 | 0.579 ± 0.077 |
| *Pinus halepensis* | 0.742 ± 0.047 | 0.688 ± 0.044 | 0.927 ± 0.027 | 0.36 ± 0.076 |
| *Pinus nigra* | 0.847 ± 0.019 | 0.768 ± 0.023 | 0.924 ± 0.017 | 0.528 ± 0.059 |
| *Pinus pinaster* | 0.765 ± 0.054 | 0.7 ± 0.057 | 0.922 ± 0.036 | 0.447 ± 0.11 |
| *Pinus pinea* | 0.765 ± 0.009 | 0.696 ± 0.009 | 0.931 ± 0.006 | 0.288 ± 0.019 |
| *Pinus sylvestris* | 0.786 ± 0.028 | 0.705 ± 0.03 | 0.973 ± 0.013 | 0.219 ± 0.04 |
| *Populus tremula* | 0.77 ± 0.028 | 0.701 ± 0.032 | 0.967 ± 0.01 | 0.197 ± 0.039 |
| *Prunus avium* | 0.804 ± 0.023 | 0.727 ± 0.022 | 0.975 ± 0.006 | 0.191 ± 0.039 |
| *Quercus ilex* | 0.767 ± 0.028 | 0.687 ± 0.029 | 0.975 ± 0.01 | 0.165 ± 0.034 |
| *Quercus petraea* | 0.789 ± 0.03 | 0.718 ± 0.032 | 0.875 ± 0.029 | 0.548 ± 0.073 |
| *Quercus pubescens* | 0.812 ± 0.017 | 0.737 ± 0.017 | 0.966 ± 0.006 | 0.241 ± 0.031 |
| *Quercus pyrenaica* | 0.778 ± 0.047 | 0.706 ± 0.047 | 0.918 ± 0.036 | 0.457 ± 0.103 |
| *Quercus robur* | 0.758 ± 0.031 | 0.696 ± 0.027 | 0.974 ± 0.009 | 0.17 ± 0.033 |
| *Salix caprea* | 0.766 ± 0.027 | 0.698 ± 0.027 | 0.936 ± 0.018 | 0.325 ± 0.049 |

### Appendix S4. Additional results using arid, mild and wet regions of the species distributions.

To calculate species-specific fixed aridity values, we derived each species’ distribution range from Caudullo et al. (2017) and constrained these ranges to the longitudinal limits of our study database. Within each constrained distribution, we generated a regularly spaced grid of points and extracted aridity values for each point. From these aridity distributions, we calculated the 15^th^, 50^th^ and 85^th^ percentiles to represent arid, mild and wet conditions, respectively, for each species. For clearer visualisation, we ultimately chose to represent only the arid and wet edges, as the mild conditions showed intermediate patterns (a full version of each figure, including mild regions, is provided in this appendix).

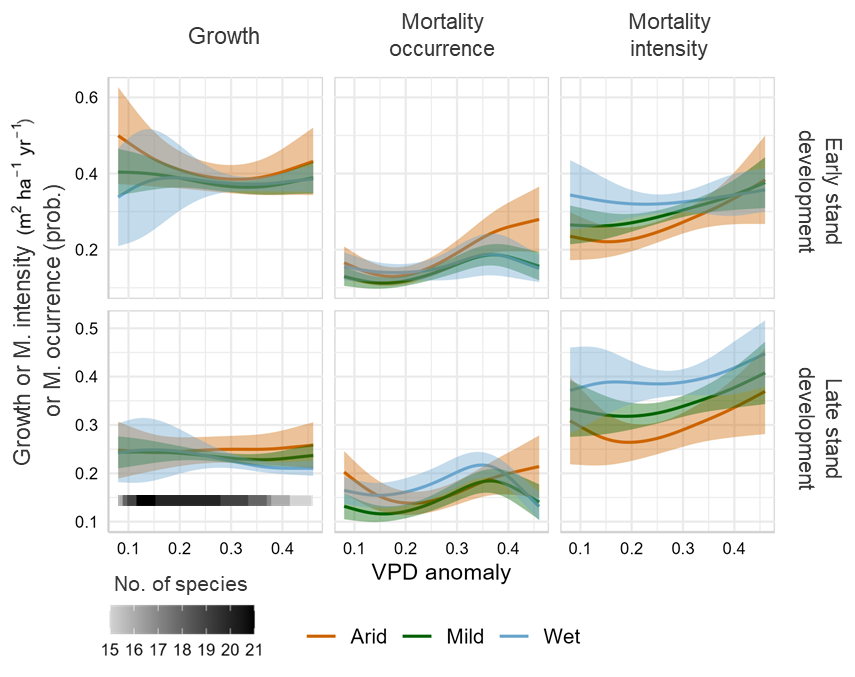

**Figure S4.1.** Weighted mean species models predictions for productivity components along the VPD anomaly gradient, restricted to the range where at least 75% of all species were present. Each species’ contribution to the mean prediction was weighted by the predictive accuracy of its model, NMAE for growth and mortality intensity models and AUC for mortality occurrence models. Shaded areas represent 95% confidence intervals derived from the weighted standard error of the species-specific predictions. The number of species contributing to the mean prediction within each segment of the VPD anomaly gradient is indicated by shading from white to black in the legend. Predictions were computed at the 25^th^ (i.e. early) and 75^th^ (i.e. late) percentiles of stand development for each species, and the species-specific arid, mild or wet edge at the 15^th^ (i.e. arid in orange), 50^th^ (i.e. mild in green) and the 85^th^ (i.e. wet in blue) percentiles of climate moisture index along the observed species’ VPD anomaly range.

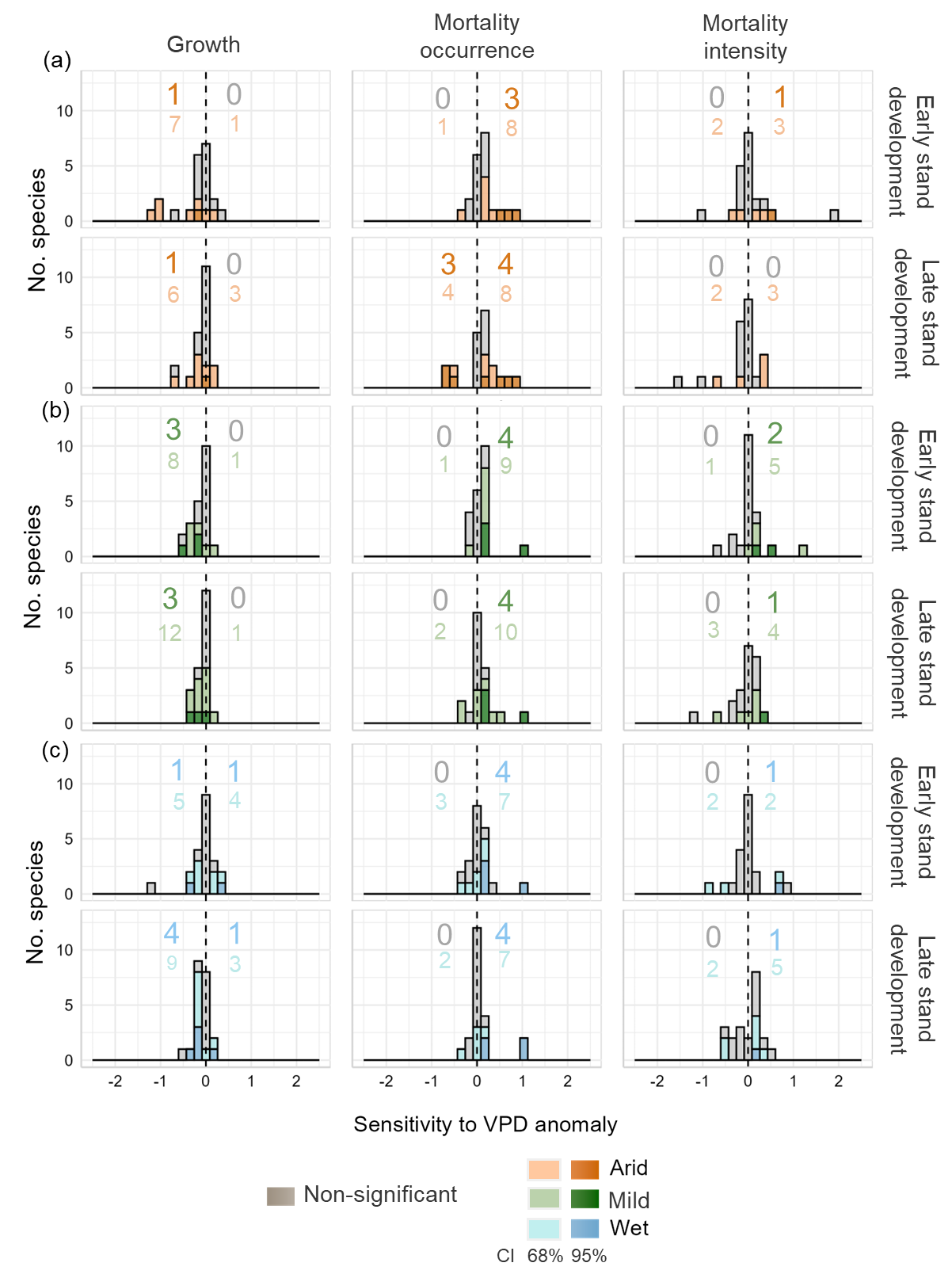

**Figure S4.2.** Species-specific sensitivity to increased VPD anomaly in (a) arid (orange), (b) mild (green) and (c) wet (blue) edges of species’ distributions and across early and late stand developmental stages. Sensitivity to VPD anomaly was calculated for each species as the difference in their response between its maximum and minimum VPD anomaly values, with bars and numbers to the right of the dashed line showing an increase in the species response with higher VPD anomaly and those to the left showing a decreased response. Dark-coloured bars and numbers represent species for which climatic sensitivity was statistically significant, with the significance being determined by whether the 95% confidence interval of the difference in the response between high vs. low VPD anomaly (±1.96 standard errors) at the species level excludes cero. Light-coloured bars and numbers indicate a less restrictive confidence interval of 68% (±1 standard errors).

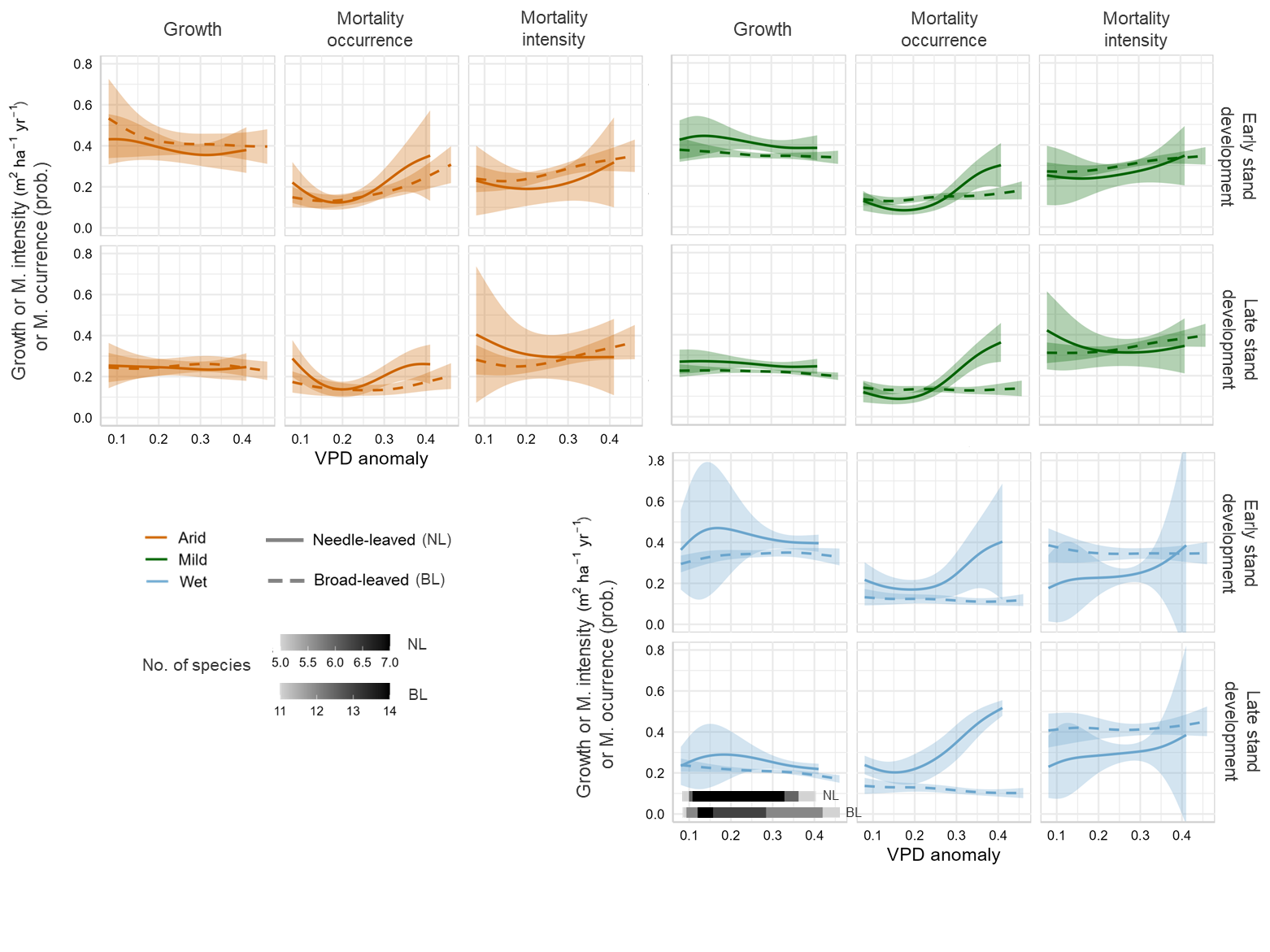
**Figure S4.4.** Weighted mean species models predictions for productivity components along the VPD anomaly gradient, restricted to the range where at least 75% of all species were present, grouped by broad-leaved and needle-leaved species. Each species’ contribution to the mean prediction was weighted by the predictive accuracy of its model, NMAE for growth and mortality intensity models and AUC for mortality occurrence models. Shaded areas represent 95% confidence intervals derived from the weighted standard error of the species-specific predictions. The number of species contributing to the mean prediction within each segment of the VPD anomaly gradient is indicated by shading from white to black in the legend. Predictions were computed at the 25^th^ (i.e. early) and 75^th^ (i.e. late) percentiles of stand development for each species, and the species-specific arid, mild or wet edge at the 15^th^ (i.e. arid in orange), 50^th^ (i.e. mild in green) and the 85^th^ (i.e. wet in blue) percentiles of climate moisture index along the observed species’ VPD anomaly range.

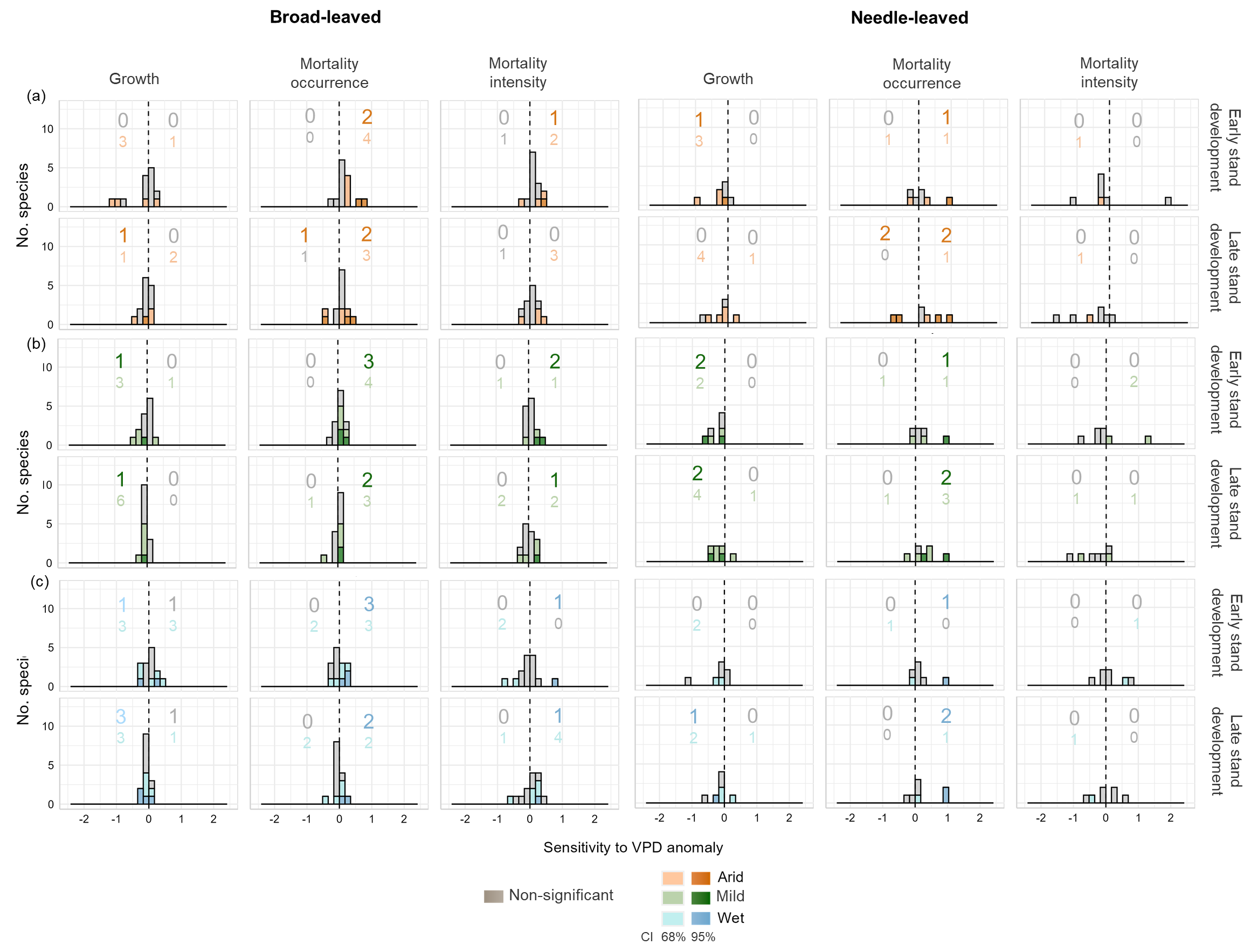

**Figure S4.5.** Broad-leaved and Needle-leaved species-specific sensitivity to increased VPD anomaly in (a) arid (orange), (b) mild (green) and (c) wet (blue) edges of species’ distributions and across early and late stand developmental stages. Sensitivity to VPD anomaly was calculated for each species as the difference in their response between its maximum and minimum VPD anomaly values, with bars and numbers to the right of the dashed line showing an increase in the species response with higher VPD anomaly and those to the left showing a decreased response. Dark-coloured bars and numbers represent species for which climatic sensitivity was statistically significant, with the significance being determined by whether the 95% confidence interval of the difference in the response between high vs. low VPD anomaly (±1.96 standard errors) at the species level excludes cero. Light-coloured bars and numbers indicate a less restrictive confidence interval of 68% (±1 standard errors).

### Appendix S5. Results at species level.

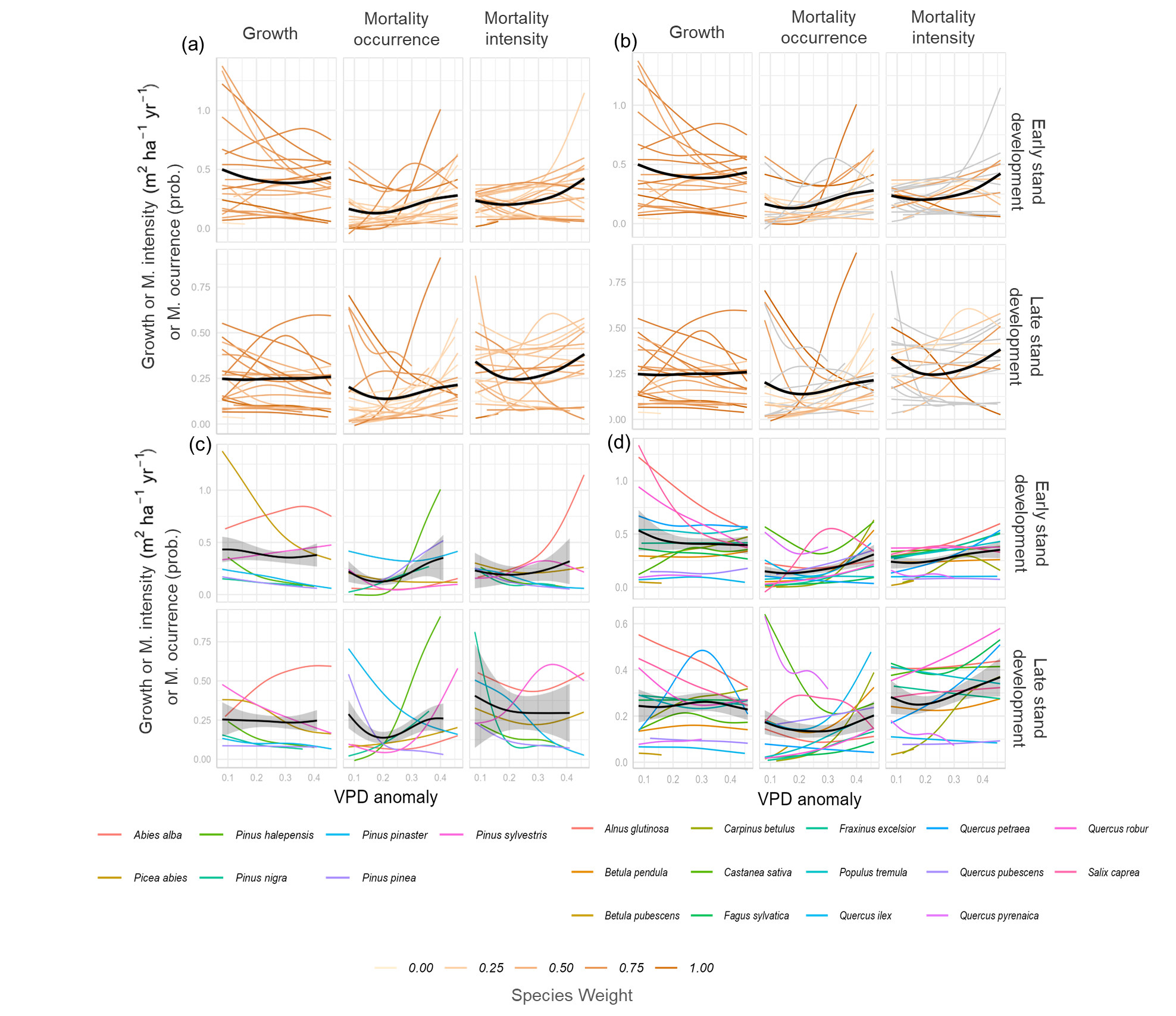

**Figure S5.1.** Species-level predictions in arid regions of the species distributions (i.e. 15^th^ percentile of species’ aridity distribution) with (a) colour representing the species-specific weight derived from the the predictive performance of its model (i.e. NMAE for growth and mortality intensity and AUC for mortality occurrence, (b) grey lines representing species for which the triple interaction was not significant in the model, (c) only needle-leaved species represented and (d) only broad-leaved species represented. Black lines represent weighted mean predictions for (a,b) all species, (c) needle-leaved and (d) broad-leaved species, with shaded areas representing 95% confidence intervals derived from the weighted standard error of the species-specific predictions.

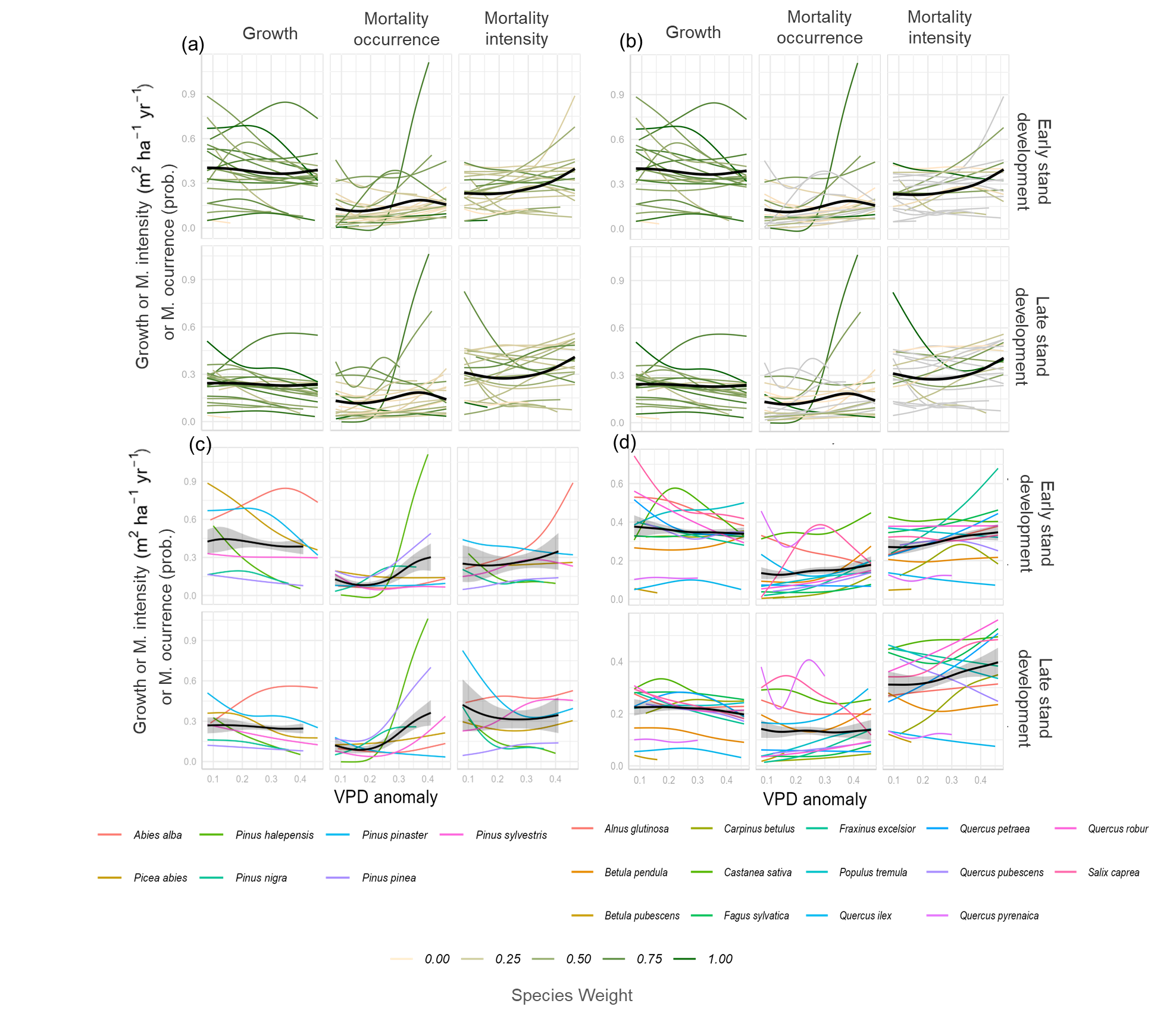

**Figure S5.2.** Species-level predictions in mild regions of the species distributions (i.e. 50^th^ percentile of species’ aridity distribution) with (a) colour representing the species-specific weight derived from the the predictive performance of its model (i.e. NMAE for growth and mortality intensity and AUC for mortality occurrence, (b) grey lines representing species for which the triple interaction was not significant in the model, (c) only needle-leaved species represented and (d) only broad-leaved species represented. Black lines represent weighted mean predictions for (a,b) all species, (c) needle-leaved and (d) broad-leaved species, with shaded areas representing 95% confidence intervals derived from the weighted standard error of the species-specific predictions.

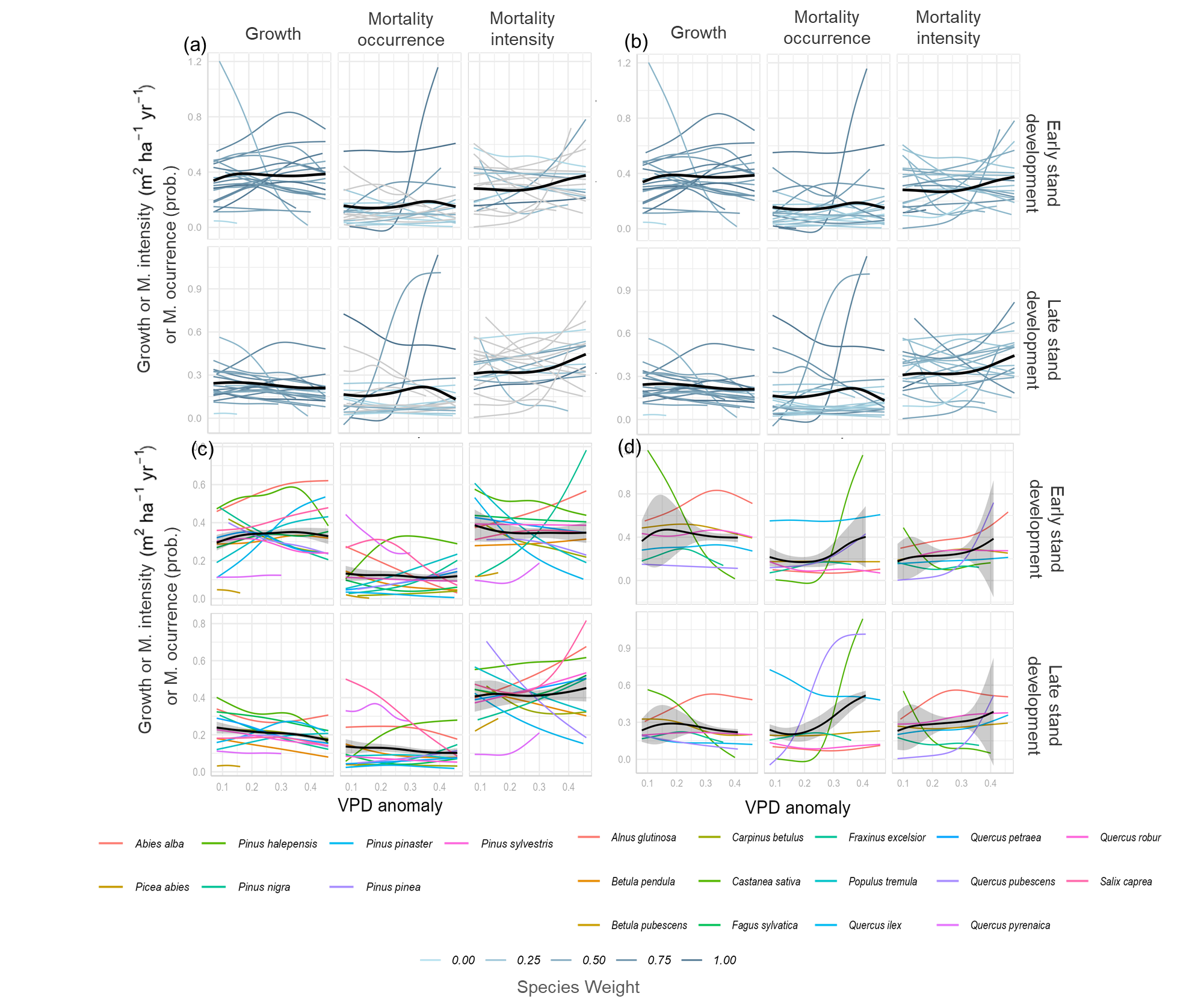

**Figure S5.3:** Species-level predictions in wet regions of the species distributions (i.e. 85^th^ percentile of species’ aridity distribution) with (a) colour representing the species-specific weight derived from the the predictive performance of its model (i.e. NMAE for growth and mortality intensity and AUC for mortality occurrence, (b) grey lines representing species for which the triple interaction was not significant in the model, (c) only needle-leaved species represented and (d) only broad-leaved species represented. Black lines represent weighted mean predictions for (a,b) all species, (c) needle-leaved and (d) broad-leaved species, with shaded areas representing 95% confidence intervals derived from the weighted standard error of the species-specific predictions.

### Figure S1. Sensitivity analysis using three alternative temperature parameters

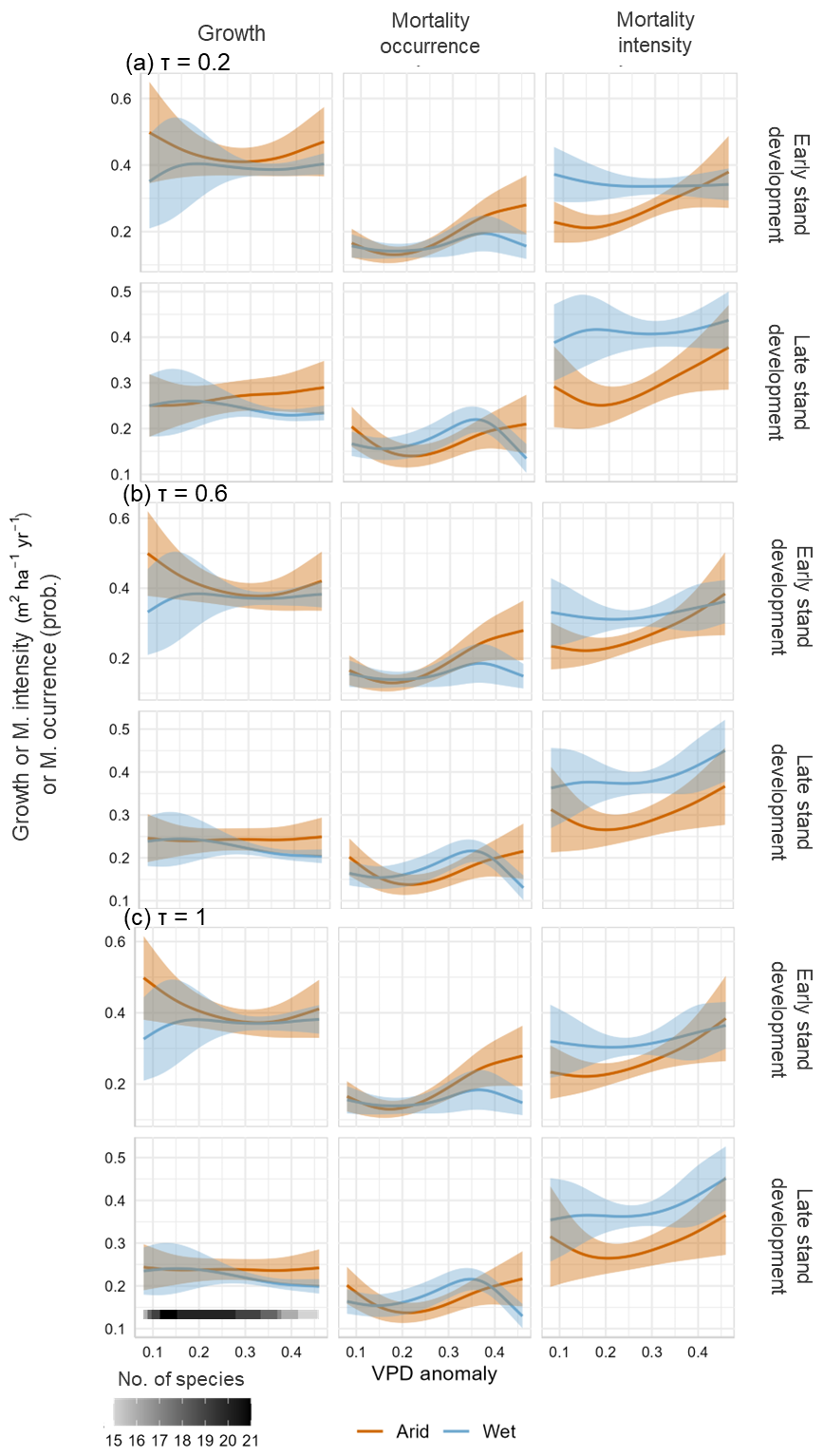

**Figure S1.** Weighted mean species predictions for productivity components along the VPD anomaly gradient, restricted to the range where at least 75% of all species were present**.** **Weights were calculated using a SoftMax function with a temperature parameter (τ) of (a) 0.2, (b) 0.6 and (c) 1.** **Smaller τ increases the differences in weight among species, while greater values allow more uniformity across species.** Each species’ contribution to the mean prediction was weighted by the predictive accuracy of its model, NMAE for growth and mortality intensity models and AUC for mortality occurrence models. Shaded areas represent 95% confidence intervals derived from the weighted standard error of the predictions. The number of species contributing to the mean prediction within each segment of the VPD anomaly gradient is indicated by shading from white to black in the legend. Predictions were computed at the 25^th^ (i.e. early) and 75^th^ (i.e. late) percentiles of stand development, and aridity edges at the 15^th^ (i.e. arid in orange), 50^th^ (i.e. mild in green) and the 85^th^ (i.e. wet in blue) percentiles of climate moisture index for each species along the observed species’ VPD anomaly range.

### Figure S2. Species-level responses to VPD anomaly (significant three-way interactions only)

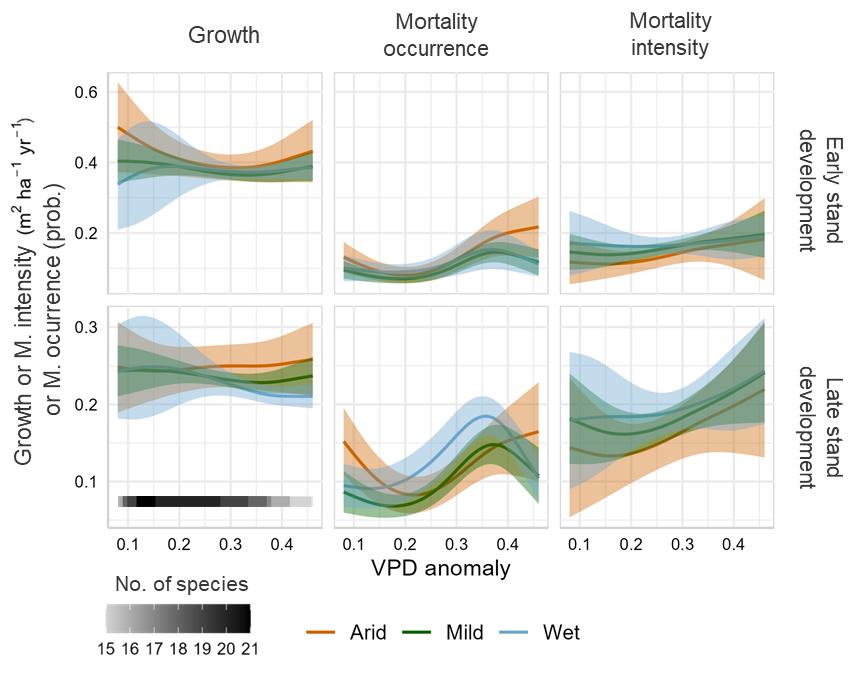

**Figure S2.** Weighted mean species predictions for productivity components along the VPD anomaly gradient, restricted to the range where at least 75% of all species were present and considering **only the species for which the three-way interaction was significant.** Each species’ contribution to the mean prediction was weighted by the predictive accuracy of its model, NMAE for growth and mortality intensity models and AUC for mortality occurrence models. Shaded areas represent 95% confidence intervals derived from the weighted standard error of the predictions. The number of species contributing to the mean prediction within each segment of the VPD anomaly gradient is indicated by shading from white to black in the legend. Predictions were computed at the 25^th^ (i.e. early) and 75^th^ (i.e. late) percentiles of stand development, and aridity edges at the 15^th^ (i.e. arid in orange), 50^th^ (i.e. mild in green) and the 85^th^ (i.e. wet in blue) percentiles of climate moisture index for each species along the observed species’ VPD anomaly range.
